# CoPhaser: generic modeling of biological cycles in scRNA-seq with context-dependent periodic manifolds

**DOI:** 10.64898/2025.12.24.696376

**Authors:** Yves Paychère, Andrea Salati, Cédric Gobet, Felix Naef

**Affiliations:** Institute of Bioengineering, Institute of Bioengineering, School of Life Sciences, Ecole Polytechnique Fédérale de Lausanne, Lausanne, CH-1015, Switzerland

**Author notes:** Contributing authors.

**Keywords:** Periodic Manifolds, scRNA-seq, Autoencoder, Biological cycles, Cell Cycle, Spatial Cancer Organization, Circadian Cycle, Menstrual Cycle, Somite Clock, Segmentation Clock, Coupling

## Abstract

Biological cycles are ubiquitous cellular processes operating across a wide range of time scales. Fundamental cycles such as the cell cycle, circadian rhythms, or the segmentation clock occur cell-autonomously and are typically coupled to other cellular processes, including cell-type identity, metabolic states, and disease-associated programs. In single-cell transcriptomics (scRNA-seq), disentangling these continuous periodic trajectories from other sources of cellular variability remains a major challenge. Here, we introduce CoPhaser, an algorithm that learns context-dependent periodic manifolds to decompose scRNA-seq count data into independent periodic and non-periodic sources of variation, while preserving interpretability of manifold coordinates across biological contexts. CoPhaser is based on a biologically informed variational autoencoder with a structured latent space that explicitly separates cycle phase from cellular context while controlling their mutual information. By modeling gene expression as context-modulated harmonic functions, the model captures flexible yet biologically grounded deformations of periodic manifolds. We demonstrate CoPhaser’s ability to yield novel biological insights across four biological cycles. It recovers accurate continuous cell-cycle phases across diverse sequencing technologies, including highly heterogeneous settings such as development and cancer, without prior knowledge of gene programs or cell-cycle states. In cancer applications, CoPhaser reveals subtype-specific proliferation dynamics, identifying quiescent primitive states in relapsed pediatric acute myeloid leukemia and distinguishing proliferation-driven from constitutive gene overexpression in triple-negative breast cancer, highlighting potential robust therapeutic targets. It further extends to spatial cancer transcriptomics, revealing spatial synchronisation of cell-cycle phases in ovarian tumors. CoPhaser generalizes to other periodic systems, enabling reconstruction of circadian clocks in the mouse aorta, and identifies cell-type and subtype-specific circadian differences. In addition, it maps continuous endometrial remodeling across the human menstrual cycle and reveals altered transcriptional dynamics in endometriosis. Finally, it reveals coupling between the cell cycle and the somite clock in the mouse embryo. Together, CoPhaser provides a versatile and interpretable framework for dissecting the interplay between cellular identity and biological cycles in single-cell data.

## 1 Introduction

The advent of single-cell genomics has accelerated the possibility of reconstructing molecularly defined biological state spaces at a fine cellular resolution [1]. An emerging property is that cells evolve dynamically along structured subspaces, commonly called manifolds, which have low dimensions compared to the high-dimensional molecular spaces [2]. Learning and representing such manifolds so that they are interpretable and biologically insightful is a major challenge, and has been successfully approached using variational autoencoders (VAEs) [3, 4], notably to predict single-cell responses to perturbations[5, 6].

To be biologically relevant, these low-dimensional spaces should respect the topological structure of the underlying processes. Indeed, these processes live in subspaces with distinct shapes, combining structures such as lines and trees, relevant in the context of cell differentiation or development, and also cycles which are common in molecular biology[4, 7].

Biological periodicity is widespread, with many important biological and cellular processes having a temporally periodic structure. Among the prime examples of biological cycles in mammals, both the cell division and circadian cycles are cell-autonomous periodic processes driven by core gene regulatory networks, including the periodic regulation of mRNA abundance through transcription and mRNA stability [8, 9]. The segmentation (somite) clock also relies on a cell-autonomous oscillatory core but additionally requires intercellular coupling and tissue-level gradients to synchronize oscillations and generate coherent traveling waves that pattern the presomitic mesoderm (PSM) [10]. These cycles span a wide range of periods: the circadian and cell division cycles operate on timescales of approximately one day, the mouse somite clock on the order of two hours, and other cycles can be substantially longer, such as the human menstrual cycle. Depending on the context, these cycles may tick only in subpopulations of cells (*e.g.* cycling tumor cells), be paused and eventually restart, or be permanently arrested.

Of particular relevance, these cycles manifest as gene expression profiles that are periodic with respect to a latent state variable, hereafter called the phase. These phases should be biologically interpretable, linking to molecularly defined events such as the G1/S transition in the cell cycle progression, the peak activity of the BMAL1/CLOCK transcription factors during the circadian cycle, or that of the *Hes7* negative regulator in the somite clock. A major challenge is to learn the phase coordinates of individual cells from (sparse) scRNA-seq counts, especially in the presence of biological heterogeneity and confounding sources of variability. Indeed, cell-autonomous cycles are embedded within cells existing in diverse biological contexts, such as different cell types, metabolic conditions, or disease progression states, or in fact any discrete or continuous cell states leading to cell state variability that cannot be explained by the cyclic process itself. Thus, while many biological cycles are often highly conserved owing to their fundamental biological functions within cells, tissues and organisms, contexts can nevertheless influence their properties, for example by shifting the means and amplitudes of the gene expression profiles (Fig. 1A). Such modified gene expression profiles complicate the assignment and interpretation of cellular phases, since then there is no fixed reference.

**Fig. 1.**
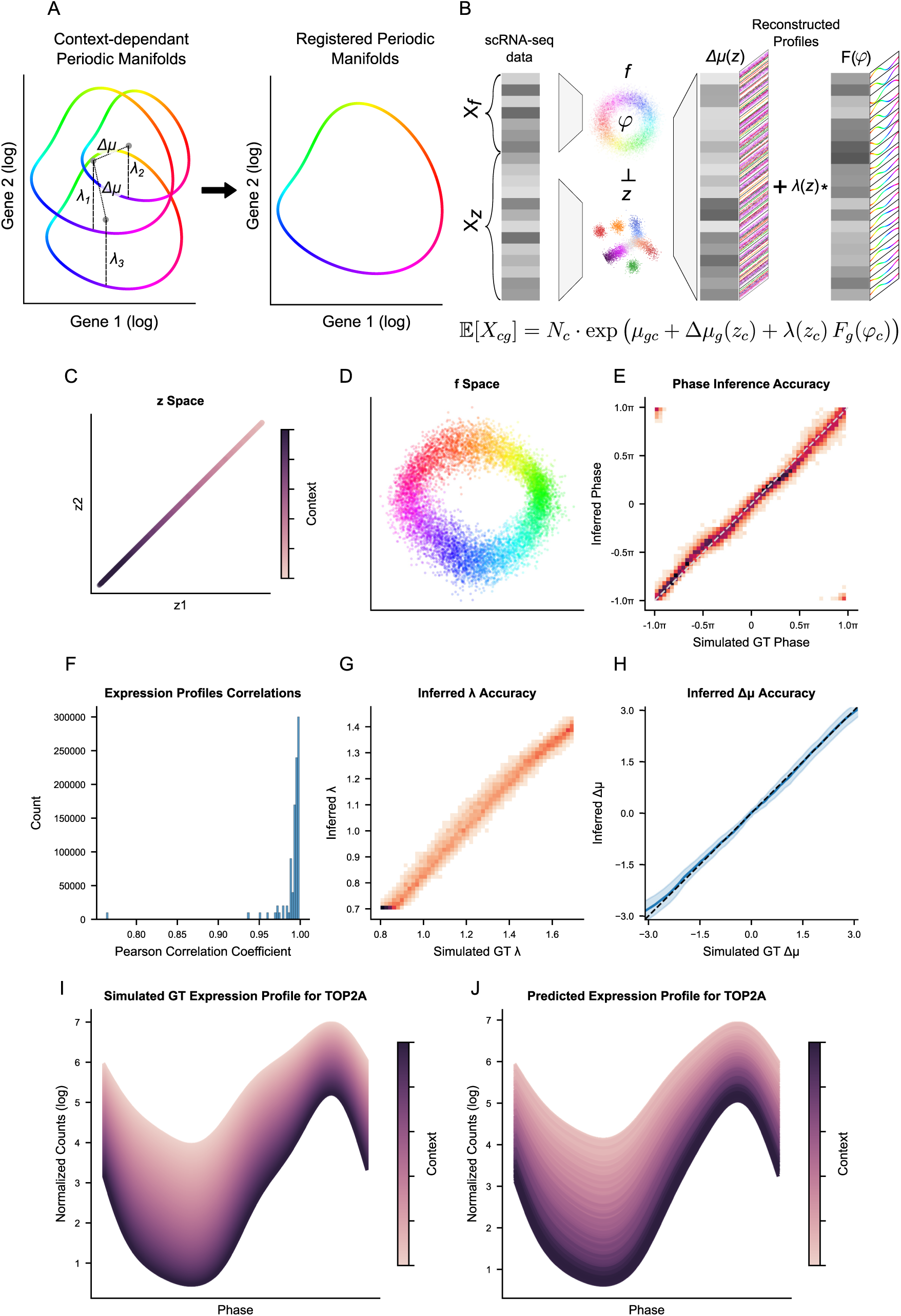
CoPhaser enables context-dependent learning of periodic manifolds using a VAE. **A.** Schematic overview of the learning task. The model addresses scenarios in which context-dependent periodic manifolds (three are shown) are registered by shifts (Δ*µ*) and scaling (*λ*) (in log). **B.** Schematic of the model architecture. Expression of rhythmic genes (*X_f_*) and context genes (*X_z_*) are projected into a two-dimensional periodic *ƒ*-space and a multidimensional contextual *x*-space, respectively. Independence between the two latent spaces is enforced by controlling their mutual information (*I*). The cyclic component of gene expression (*Fg*) is reconstructed from the cell-specific phase (*φ*) in *ƒ*-space using Fourier series. Gene- and cell-specific mean shifts (Δ*µg* (*zc*)) and cell-specific rhythmic amplitudes (*λ*(*zc*)) are decoded from *x*-space. Expected gene counts (E[*Xcg*]) are obtained by combining the library size (*Nc*) with the inferred mean expression fraction (*µcg*). **C.** *x*-space inferred from simulated data containing a one-dimensional contextual variable. **D.** Inferred *ƒ*-space from simulated data, colored by the simulated ground-truth (GT) phase. **E.** Scatterplot comparing inferred and simulated GT phases, colored by local density (darker indicates high density). **F.** Pearson correlation coefficient between inferred and simulated GT gene expression profiles evaluated at 20 evenly spaced phases in [*−π, π*) using inferred and simulated GT Fourier coefficients for a fixed context (Methods ^4^.^7^). **G.** Scatterplot comparing inferred and simulated GT rhythmic amplitudes (*λ*(*zc*)), colored by the local density. **H.** 95 % confidence intervals of inferred mean shifts across the range of simulated GT values across all genes and contexts. **I.** Simulated expression profile of DNA topoisomerase II alpha (*TOP2A*) across the one-dimensional context. Realistic Fourier coefficients for the simulations were obtained from fitting RPE1 scRNA-seq data [29], and Δ*µ* and *λ* were randomly generated as functions of the contextual variable. **J.** Reconstructed *TOP2A* expression profile using the inferred Fourier coefficients, Δ*µ*, and *λ*.

The cell cycle is ubiquitous in development and regeneration, while it is also frequently mis-regulated in disease. Numerous methods have sought to estimate cell cycle states from scRNA-seq (reviewed in [11]). Such methods typically seek to either regress out cell-cycle related variability for downstream analyses [4, 12], or study gene expression dynamics [13]. Common models for cell cycle phase inference consist of autoencoders (AEs), of which Cyclum [14] and DeepCycle [15] are prominent representatives. These AEs infer a latent phase describing cell-cycle progression and enables high-resolution reconstruction of transcriptional trajectories in deeply sequenced, actively proliferating, and homogeneous systems. While successful in homogeneous populations of cells, these methods do not consider that the cell-cycle can be modified by the cellular context, limiting their applicability in heterogeneous situations such as development [16] or disease [17, 18].

While the abundance and detectability of cell cycle transcripts in mammalian cells is favorable for the phase inference task, other cycles are more challenging as the availability of sufficiently highly expressed periodic transcripts can be more scarce. This is notably the case for the circadian cycle, where phase assignment algorithms have been most successful in bulk [19, 20]. Nevertheless, the assignment of circadian phases in single cells is possible when the sequencing coverage is sufficient [21, 22]. In particular, Tempo [21] implements a probabilistic model for Bayesian circadian phase inference in a single context that uses variational inference. By modeling rhythmic gene expression using shared oscillatory waveforms across cells, Tempo can allocate phase coordinates in homogeneous circadian systems. However, Tempo does not explicitly model cell-type or state-dependent modulation of circadian gene expression. How to approach circadian heterogeneity and contextual dependence at the single-cell level is still open, and can be approached in multiple ways, including the recent VAE VIST [23]. VIST generically decomposes scRNA-seq time-series data into time-dependent and time-independent components, facilitating joint batch correction and temporal analysis. VIST is designed for arbitrary time-series designs and hence does not impose an explicit periodic topology on the latent space, and requires observed experimental time, making it well-suited to synchronized or externally driven time courses. Similarly, CellUntangler [24] is a VAE designed to disentangle different biological processes, including cycles. However, it does not register cells on a strict one-dimensional cyclic manifold, and its architecture is not designed to yield a phase definition that is comparable across different contexts.

Here, we formalize the idea that context-dependent periodic manifolds can be learned in an unsupervised manner, enabling the definition of phase coordinates that remain interpretable and comparable across cellular contexts. We develop a generic model, *CoPhaser* (for <u>co</u>ntext-aware phaser), applicable to biological cycles and data modalities, without requiring external labels or prior knowledge of periodic gene profiles, while remaining flexible enough to support transfer learning from independently learned or calibrated cycles. With its flexible but explicit parameterization of the expected expression levels in function of the latent phase and context coordinates, CoPhaser consists of a biologically informed variational autoencoder with a structured latent space that explicitly separates cycle phases from cellular context, while controlling their mutual information. By modeling gene expression as context-modulated harmonic functions, CoPhaser captures flexible deformations of biological cycles.

We demonstrate that CoPhaser recovers biologically accurate continuous cell-cycle phases across diverse RNA sequencing technologies and highly heterogeneous situations such as development and cancer, without requiring prior knowledge of gene expression programs or cell-cycle states. We applied CoPhaser to three cancer datasets, revealing subtype-specific proliferation dynamics, identifying quiescent primitive states in relapsed pediatric acute myeloid leukemia, and distinguishing proliferation-driven gene overexpression from constitutive overexpression to uncover robust therapeutic targets in triple-negative breast cancer. Thirdly, CoPhaser seamlessly handles spatially resolved cancer transcriptomics, showing that proliferative states are not stochastically distributed but instead form spatially coherent domains within the tumor architecture. By mapping continuous phases onto tissue coordinates, we identify localized niches of synchronized cell-cycle progression and demonstrate that even within morphologically homogeneous tumor regions, cellular proliferation is organized into distinct, phase-stratified neighborhoods. Furthermore, CoPhaser generalizes well to other periodic systems, enabling the reconstruction of circadian clocks in the mouse aorta and the identification of cell-type– and subtype-specific circadian differences. We then employed CoPhaser to map continuous endometrial remodeling across the human menstrual cycle, uncovering distinct transcriptional dynamics in women with endometriosis compared with controls. Finally, we applied CoPhaser to reconstruct the somite clock along the anterior-posterior axis in the mouse PSM, where it revealed traveling waves from snapshot transcriptomic data and a coupling between the cell cycle and the segmentation clock.

## 2 Results

### 2.1 CoPhaser enables context-dependent learning of periodic manifolds

While many biological cycles are highly conserved, they are rarely fully invariant. Instead, their gene expression trajectories measured in single-cell RNA-sequencing (scRNA-seq) manifest as periodic manifolds that can be influenced by the underlying cellular context (Fig. 1A). To register such cycles across contexts, we allowed the manifolds to be deformed by shifting and scaling according to a context variable. Specifically, to disentangle these periodic dynamics from cell-specific context, we developed CoPhaser, a variational autoencoder (VAE) with a structured latent space composed of an *f*-space encoding a periodic phase and a cell-context *z*-space (Fig. 1B). The model uses factorized variational posteriors consisting of a Power Spherical distribution [25] for the phase and a multivariate Gaussian for the non-periodic latent variable *z*. The decoders, yielding the distributions of the measured scRNA-seq counts for each cell, consist of Fourier series to capture the periodic structure and standard decoders for the context-dependent mean and amplitude corrections (Fig. 1B and Methods 4.1).

The encoder is informed with two different sets of genes: a set of seed rhythmic genes (*X_f_*) determines the *f*-space, whereas the *z*-space is inferred from a broader set that includes highly variable genes (Methods 4.8.1). To ensure the identifiability of the *f* and *z* coordinates and avoid that cyclic variability is attributed to the more general *z*-space, we simultaneously train a Mutual Information Neural Estimator (MINE) [26] to explicitly penalize dependencies between the two latent spaces (Methods 4.3). Formally, for each cell *c* and gene *g*, the log-expected gene counts log E[*X_cg_*] are modeled as the sum of three contributions: a periodic component *F_g_*(*φ_c_*), a baseline expression *µ_cg_*, and a context-dependent mean shift Δ*µ_g_*(*z_c_*). In addition, we allow for a context-dependent cycling amplitude *λ*(*z_c_*) that scales the periodic component *F_g_*(*φ_c_*) (Fig. 1B). This parameterization enables the model to learn flexible, context-dependent deformations of the periodic manifold, while preserving a coherent and shared definition of the phase (*φ_c_*) across cells. The model is trained using a custom loss based on the Evidence Lower Bound (ELBO), with observation noise modeled by a Negative Binomial distribution (Methods 4.4), augmented with several loss terms to favor biological solutions (Methods 4.4).

### 2.2 Accurate recovery of phases and context-dependent modulations in simulated data

To validate our approach, we simulated data recapitulating cell-cycle gene expression counts using biologically plausible Fourier coefficients (Methods 4.7). We also introduced a one-dimensional cellular context that linearly modulates both the mean gene expression levels and the cycle amplitude. CoPhaser correctly decomposed these sources of variations: the inferred *z*-space accurately recovers the contextual gradient (Fig. 1C), while the *f*-space faithfully encodes the ground-truth (GT) cell-cycle phase (Fig. 1D,E). Consequently, the inferred gene expression profiles (Fig. 1F), cycle amplitudes (Fig. 1G), and mean shifts Δ*µ* closely match the simulated GT, resulting in accurate reconstruction of gene expression even in the presence of strong contextual distortions (Fig. 1I,J). To assess the roles of our model components, we performed ablation studies. We found that removing the contextual *z*-space resulted in a model that fails to distinguish between cyclic and contextual variance, producing severe artifacts in the inferred phases (Fig. S1A-C). This limitation is also observed in standard Principal Component Analysis (PCA), which similarly lacks the capacity to model context-dependent deformations and thus confounds the two sources of variation (Fig. S1D-F). Furthermore, controlling the mutual information (*I*) between the *f* - and *z*-spaces proved to be critical, as removing this constraint led to information leakage and degraded separation of the two latent representations (Fig. S1G–K).

### 2.3 Context-aware inference of continuous cell-cycle phases across datasets and technologies

We validated CoPhaser on cell-cycle analysis in scRNA-seq datasets of increasing complexity, using a literature-based cycling gene set comprising 98 genes (Methods 4.8). First, using scEU-seq data from the human RPE1-FUCCI cell line [8], we observed a monotonic relationship between the inferred phase and the FUCCI-derived cell cycle phases (Fig. 2A). Note that this monotonic, but non-identical, relationship between FUCCI- and mRNA-derived phases is expected as the two methods are reporting on biochemical processes occurring on different regulatory layers. In fact, the FUCCI-based phase appears less informative in the interval ((inferred phase ∈ (−0.5*π,* 0)), resulting in an apparent compression and discontinuity in gene expression profiles in late G1 (FUCCI phase ∼ −0.5*π*), whereas CoPhaser infers a smooth progression, indicating that mRNA contains biologically relevant information on cell-cycle progression when the FUCCI signal is not sensitive (Fig. 2B). The latent context *z*-space captures the experimental conditions in the original study (Fig. S2A), however, in this case, without inducing substantial differences in the predicted gene profiles (Fig. S2B). This is consistent with the high correlation between CoPhaser and PCA-based phase inference on this dataset (Fig. S2C,D). Indeed, PCA-based phases are reliable when the context-induced variability is limited [27]. Moreover, all methods capture the expected monotonic increase in the mean number of unique molecular identifiers (UMIs) per cell over the cycle (Fig. S2D). Finally, the inferred peak phases of cell-cycle genes show a good agreement with the discrete independent annotations (G1, S, G2/M) from Cyclebase [28] (Fig. S2E).

**Fig. 2.**
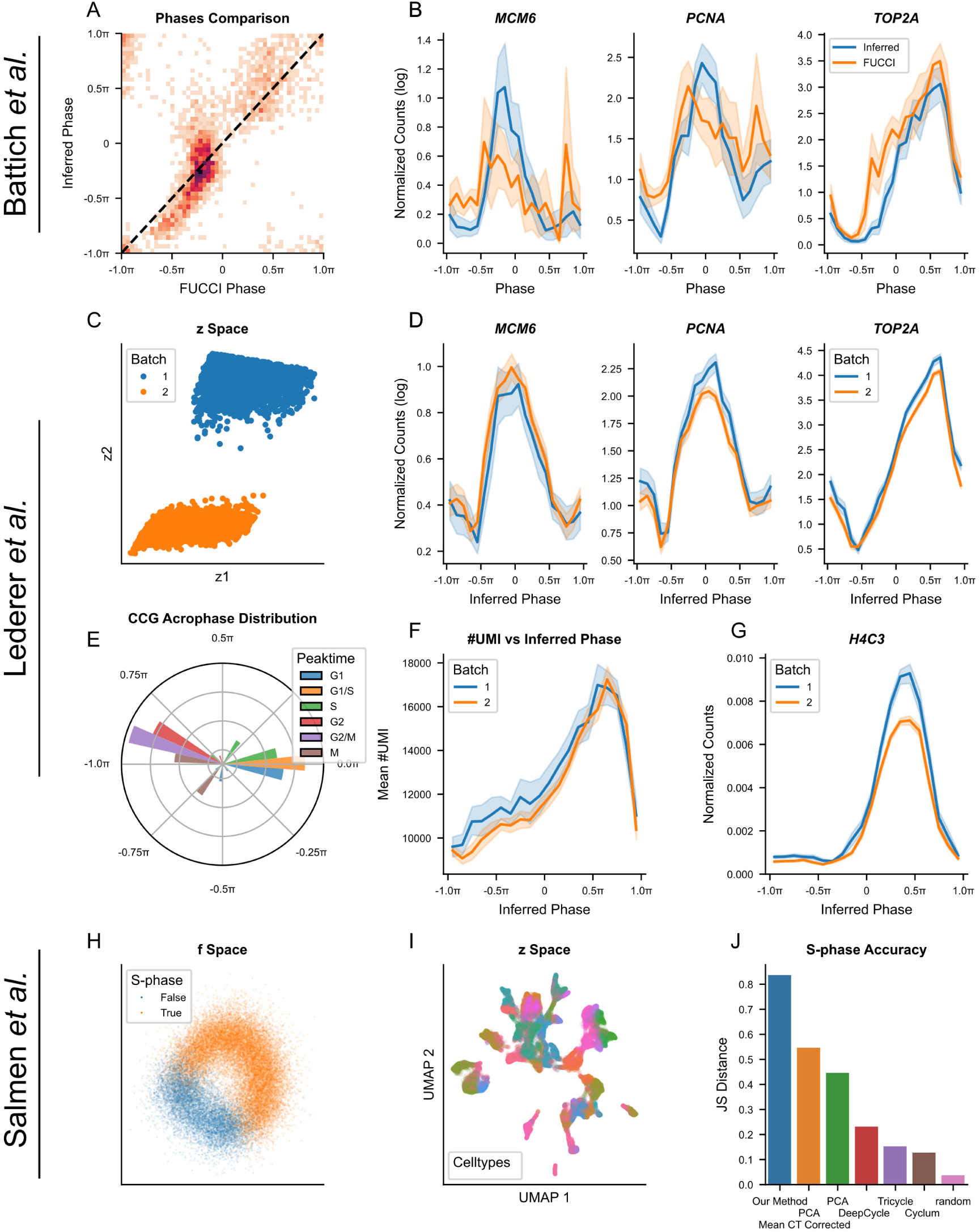
Context-aware inference of continuous cell-cycle phase across datasets and technologies. **A,B.** Cell-cycle phase inference on RPE1-FUCCI scRNA-seq data from [8]. **A** Comparison between inferred phases and FUCCI-derived ground-truth phases, color coded by local density. **B** Reconstructed gene expression profiles using inferred phases (blue) and FUCCI-derived phases (orange). Profiles are obtained by binning cells by phase and averaging log(1 + counts per 10,000) for the indicated gene. **C–G.** Cell-cycle phase inference on two RPE1 scRNA-seq batches (10x Genomics) from [29]. **C,** Inferred *x*-space colored by batch identity. **D** Reconstructed gene expression profiles colored by batch, obtained by binning cells according to inferred phase and averaging log(1 +counts per 10,000) for the indicated gene. **E.** Distribution of inferred peak phases for cell-cycle genes, colored by their annotated peak phase according to Cyclebase [28]. **F.** Mean number of unique molecular identifiers (UMIs) per cell as a function of inferred phase, colored by batch. **G.** Mean fraction of counts derived from histone genes as a function of inferred phase, colored by batch. **H–J.** Cell-cycle phase inference on mouse embryonic scRNA-seq data (VASA-seq) from [16]. **H.** Projected *ƒ*-space colored by the original S-phase annotations [16]. **I.** Projected *x*-space using a Uniform Manifold Approximation and Projection (UMAP) colored by original cell-type annotations [16]. **J.** Jensen–Shannon divergence (base-2 logarithm) between the original S-phase annotation and binned inferred phases, computed for CoPhaser, principal component-based baselines (PC1 and PC3 yielded the best score; mean-centered either by cell type or across all cells), DeepCycle [15], Tricycle [50], Cyclops [19], and a random phase assignment.

We further evaluated the robustness of the latent representations using two batches of RPE1 scRNA-seq data (10x Genomics) [29]. While these experiments were performed in the same conditions, the *z*-space was sensitive enough to separate the two batches (Fig. 2C). The inferred phases recapitulate canonical cell cycle gene expression patterns (Fig. 2D) with the correct temporal ordering of peak phases (Fig. 2E). Consistent with prior knowledge, the total UMI count increases monotonically across the cycle (Fig. 2F) and sharply drops following mitosis. Furthermore, the histone transcript H4C3, which is not part of the rhythmic input gene set, consistently shows a sharp increase followed by a rapid decrease bracketing the inferred S-phase alongside known markers, providing an independent validation of the high resolution of the inferred phases (Fig. 2G).

A potential limitation of our approach is the requirement for an input list of cycling genes. However, because Fourier series are also inferred for all context input genes, we hypothesized that this list could be refined iteratively for cycles that induce high amplitude gene expression signatures. We tested this strategy on the RPE1 dataset by initializing the model with a minimal set of just two cycling genes, *TOP2A* and *PCNA*, while using the 2000 most variable genes as context genes. This minimal setup already yielded a good approximation of the cell cycle, with a circular correlation of 0.47 relative to the reference (Fig. S3A, B). We note however that despite the inferred gene peaks following the correct temporal ordering (Fig. S3C), the inferred amplitude of *PCNA* became exaggerated (Fig. S3B), indicating overfitting, particularly in the low-expression interval (−*π,* 0). This overfitting manifested in artifacts in both the reconstructed UMI counts and histone expression profiles (Fig. S3D,E).

To test how performance improves with more cycling genes, we defined a gene selection score combining mean expression and inferred amplitude to identify robust markers (Methods 4.8.1). We selected the twenty most informative genes and retrained the model. Using this refined gene set, the inferred phases closely match those obtained with the literature-based 98-gene list (Methods 4.8), without visible artifacts and faithfully recapitulating the expected biological cell cycle gene expression dynamics (Fig. S3F-J).

Finally, we evaluated our method on a more complex dataset of mouse embryonic cells profiled by VASA-seq [16], which was provided with independent S-phase annotations. Despite the rich cellular heterogeneity and numerous proliferative cell types present in the embryos, the inferred *f*-space cleanly separates S-phase cells from other populations (Fig. 2H). As expected, the dominant source of variation in the *z*-space corresponded to cell identity (Fig. 2I), closely resembling the original embedding [16]. Crucially, owing to the explicit separation of latent spaces, the *z*-space is largely devoid of cell cycle–related variation (Fig. S4A). Histone expression increases and peaks during S-phase, followed by a decline as G2 genes peak, in agreement with established cell cycle biology (Fig. S4B, C). However, significant shifts in the context-dependent means are visible between cell types, with genes peaking in G1 or at the G1 to S transition being more affected than the others (Fig. S4D). As prominent examples, *Cdca7*, which peaks at the S-phase transition, displays large mean shifts between cell types, while *Kif11*, peaking in G2, displays none. To systematically assess performance, we computed the Jensen–Shannon (JS) divergence between the inferred phase distributions of annotated S-phase cells and non–S-phase cells. A perfect separation yields a score of 1, whereas random assignments yield 0. Here, CoPhaser outperformed all alternatives, including PCA projections, whether mean-centered per cell type or globally (Fig. 2J). The assignments from Seurat perform poorly, with a separation close to random (Fig. S4G), and the leading PCs yield only moderate separation (Fig. S4H). In contrast, our approach achieves a clear separation with limited overlap confined to cell cycle transitions (Fig. S4I), underscoring the advantages of a structured latent space and explicit modeling of context-dependent cycle deformations.

### 2.4 Analysis of subtype-specific proliferation rates and phase-dependent versus constitutive gene dysregulation in breast cancer

Having validated our model on controlled and heterogeneous non-tumoral datasets, we next investigated its performance in highly variable tumoral contexts. Tumors often contain a large fraction of quiescent cells in G0/G1, which can obscure cell cycle inference. To address this, we augmented our model with a simultaneously trained Bernoulli VAE [30] that generically classifies cells as cycling or non-cycling, thereby reducing the influence of quiescent cells on phase inference (Methods 4.2). Specifically, the Bernoulli encoder takes as inputs all cyclic genes and the latent context *z*, and the learned indicator functions then multiply the Fourier series so that cells classified as non-cycling are not influenced by the periodic gene expression.

We first analyzed a breast cancer dataset comprising 69 surgical tissue specimens from 55 patients [31], focusing exclusively on tumoral epithelial cells. We used this dataset, for which the original authors provided a cycling annotation, to validate our cycling classifier. Cells identified as non-cycling by the Bernoulli encoder are indeed restricted to the G0/G1 interval (Fig. S5A). Our classification was less strict than the original annotation (Fig. S5B, C); this difference likely reflects that the original annotation relies on *Mki67* mRNA expression, which is very low during G1, potentially leading to false negatives in the original annotation. Feature attribution via Integrated Gradients [32] between cells in G1 and cells classified as non-cycling, displayed shared negative markers between subtypes consisting mostly of cell cycle regulators, as well as fewer positive markers, among which epithelial markers such as keratins (Fig. S5D–F).

Analyzing the predicted cycling logits for cells inferred to be in G1 or G0, we observed a striking divergence between breast cancer subtypes, with triple-negative (TN) tumors assigned with a markedly higher probability of cycling (Fig. S5). To further characterize the signal driving this classification, we compared the gene expression signatures of G0/G1 cells classified as cycling and non-cycling. This analysis revealed an upregulation of cell cycle–associated genes, most notably the proliferation marker *BIRC5* (Fig. S5H, I), suggesting that the classifier identifies a G0-like state. Having validated our classifier, we investigated the predictions of the other components of our model. In this dataset, the local structure of the *z*-space is dominated by inter-patient variability (Fig. S6A), while the global organization preserves tumor subtypes (Fig. 3A). As observed previously, the inferred peaking times of cell cycle genes follow the expected order (Fig. 3B), and *H4C3* peaks with S-phase genes, exhibiting similar profiles across subtypes (Fig. 3C). Gene expression profiles are well aligned between subtypes (Fig. S6B), and the mean UMI count increases monotonically along the inferred cell cycle (Fig. S6C).

**Fig. 3.**
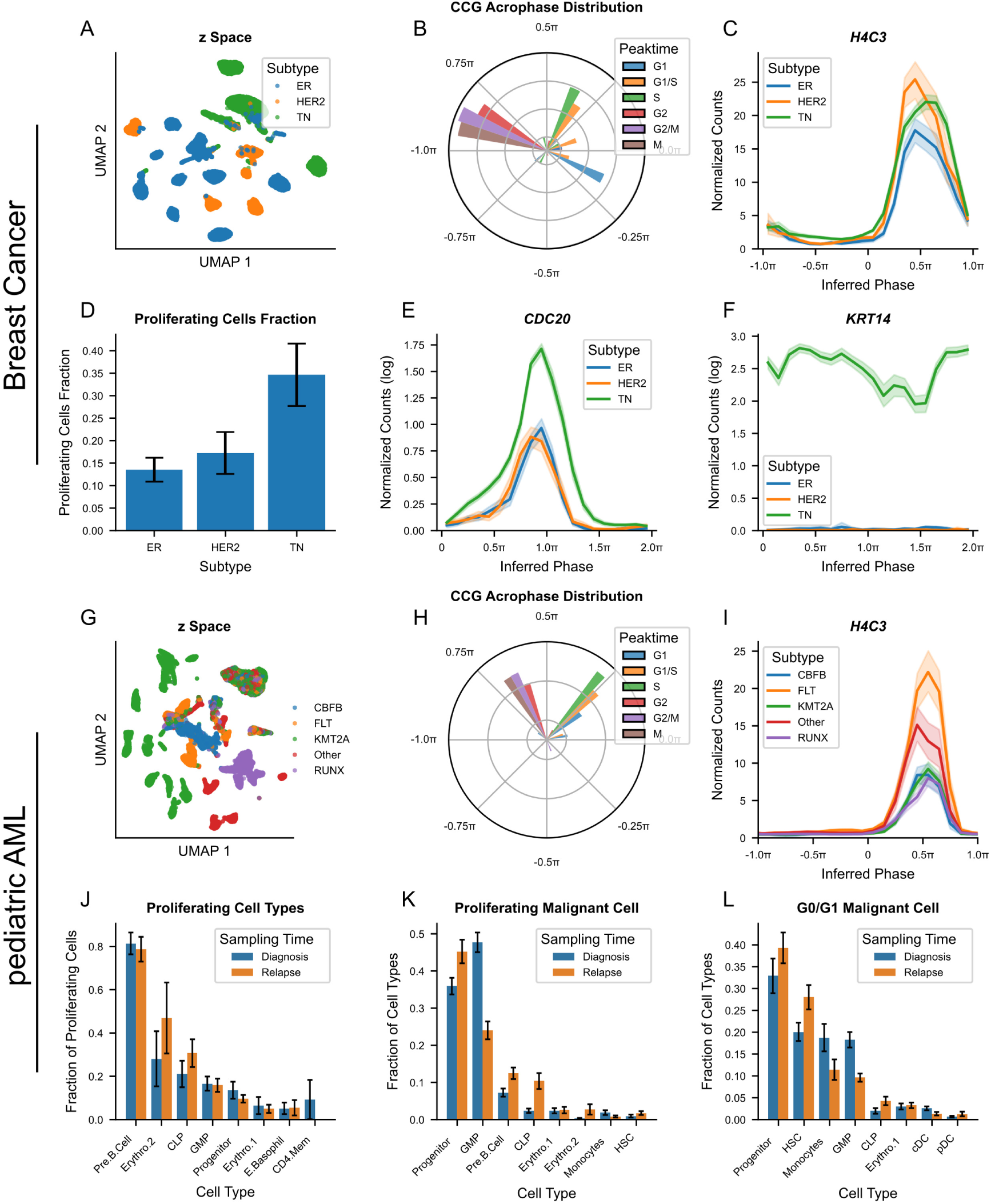
CoPhaser reveals subtype-specific proliferation patterns in breast cancer and AML. **A–F.** Analysis of breast cancer scRNA-seq data from [31]. **A.** UMAP projection of the inferred contextual *x*-space colored by clinical subtype (ER: Estrogen receptor positive; HER2: human epidermal growth factor receptor 2 positive; and TN: triple-negative) **B.** Polar histogram of inferred peak phases for cell-cycle genes, colored by their annotated peak phase according to Cyclebase [28]. **C.** Reconstructed expression profile of *H4Ch* as a function of inferred phase, colored by clinical subtype. **D.** Fraction of proliferating cells across tumor subtypes. **E,F.** Reconstructed expression profiles of *CDC20* (**E**) and *KRT14* (**F**) as a function of inferred phase, colored by clinical subtype. Profiles were obtained by binning cells according to their inferred phase and averaging log-normalized expression (log(1 + counts per 10,000)). **G–L.** Analysis of pediatric acute myeloid leukemia (AML) scRNA-seq data from [35] **G.** UMAP projection of the inferred contextual *x*-space, colored by tumor subtype. **H.** Polar histogram of inferred peak phases for cell-cycle genes, colored by their annotated peak phase according to Cyclebase [28]. **I.** Reconstructed expression profile of *H4Ch* as a function of the inferred phase. Profiles were obtained by binning cells according to their inferred phase and averaging normalized expression (counts per 10,000) for each tumor subtype. **J.** Fraction of proliferating malignant cells for each cell type in diagnosis and relapse patients. Erythro.1: early erythrocytes; Erythro.2: late erythrocytes. Error bars represent the standard error of the mean (SEM) across patients. **K,L.** Cell-type composition of the proliferating malignant cells (**K**) and G0/G1 malignant cells (**L**). Error bars represent the SEM across patients.

By quantifying the fraction of proliferating tumoral cells, we identified significant differences between subtypes, with triple-negative breast cancer (TN) exhibiting the highest proliferation rates (Fig. 3D). Analysis of inferred mean shifts further revealed that TN cells display elevated expression of *CDC20* (Fig. 3E), consistent with prior reports [33]. However, since *CDC20* is not constitutively overexpressed but rather upregulated specifically during the S/G2/M phases, the predicted expression dynamics suggest limited efficacy of therapies targeting *CDC20* in slow-dividing populations such as persistent cancer cells [17]. In contrast, CoPhaser identifies genes such as *KRT14* as being constitutively overexpressed in TN cells (Fig. 3F), consistent with their role as known markers of TN breast cancer [34].

An advantage of our model is that the inferred mean shifts are decoupled from differences in proliferation rates. For example, pseudobulk expression of *UBE2C* suggests higher expression in TN tumors (Fig. S6D). However, inspection of its expression profile along the cell cycle (Fig. S6E) reveals dynamics similar to other subtypes, indicating that the apparent overexpression arises primarily from the increased fraction of cycling cells rather than a true context-specific upregulation.

### 2.5 CoPhaser reveals persistence-driven relapse in pediatric AML

We next analyzed a longitudinal pediatric acute myeloid leukemia (pAML) dataset comprising scRNA-seq data from 28 patients sampled at diagnosis, remission, and relapse [35]. In this dataset, the contextual *z*-space was primarily structured by tumor subtype and patient ID (Fig. 3G, Fig. S6F), with non-tumoral cells clustering together (Fig. S6G). This organization highlights the dominant variability induced by tumor driver mutations, which exceeds that associated with the diverse cell types present in AML (Fig. S6H). Despite this heterogeneity, the inferred cell cycle is robust: the peaking times of canonical genes follow the expected temporal progression, with a notably prolonged G1 phase (Fig. 3H), and the histone gene *H4C3* consistently peaking in S-phase genes across all subtypes (Fig. 3I). Moreover, gene expression profiles are well aligned across subtypes, differing mainly by context-dependent shifts in the (log) means (Fig. S6I), and the mean number of UMIs per cell increases monotonically over the inferred cell cycle (Fig. S6J).

We next quantified the fraction of actively proliferating malignant cells, defined as cells with an inferred phase in (*π/*4*, π*) and predicted to cycle, corresponding to non-G0/G1 cells, for each malignant cell type. Pre-B cells, late erythrocytes, common lymphoid progenitors (CLP), granulocyte–monocyte progenitors (GMP), and progenitor cells exhibited the highest proliferative fractions (Fig. 3J). Notably, the fraction of cycling cells within each cell type did not change significantly between diagnosis and relapse across patients. In contrast, examining the composition of proliferating malignant cells revealed a marked relative decrease in GMP cells and a concomitant increase in progenitor cells at relapse (Fig. 3K). Analysis of non-cycling (G0/G1) malignant/tumor cells further confirmed a global shift toward more primitive cell states following treatment (Fig. 3L), consistent with the original model [35]. Importantly, this shift appears to be driven by persistent cancer cells, displaying a low proliferation rate [17] and low cell death, as these primitive populations do not cycle more frequently (Fig. 3D). Hematopoietic stem cells (HSCs) exemplify this behavior: although rarely proliferating, they constitute a substantial fraction of G0/G1 cells, reflecting their long life time.

### 2.6 Spatial mapping of cell-cycle phases identifies clusters of synchronized cell-cycle states in ovarian cancer tissues

While cell proliferation is a defining characteristic of cancer, its spatial topography in solid tumors is still poorly characterized. Prior work across multiple cancer types suggested that proliferation status can be organized into distinct physical domains over multiple spatial scales, ranging from small cellular niches to larger structured neighborhoods [18]. Traditional assessments using single protein markers like Ki-67 frequently fail to capture the continuous nature of cell cycle dynamics and the specific niches that foster proliferative or arrested states [36].

Here, we applied CoPhaser to spatial transcriptomics (Xenium Prime 5K) to investigate the spatial topography of cell-cycle phases within human ovarian adenocarcinoma. We analyzed two samples with contrasting clinical and technical profiles: a metastatic High-Grade Serous Carcinoma (HGSC) preserved as Formalin-Fixed Paraffin-Embedded (FFPE) [37] and a second adenocarcinoma sample preserved as Fresh Frozen (10x Genomics). To ensure cross-sample comparability, we performed joint inference of the phase and *z*-space across both samples. This joint fitting of approximately 1.5 million cells leverages Cophaser’s ability to ensure that the recovered cell-cycle phases and underlying latent topography are directly comparable between the datasets (Fig. S7 A). Despite technical differences (sample preservation, coverage), CoPhaser successfully resolved the cellular heterogeneity in both samples. The integrated contextual *z*-space identified three distinct tumor epithelial clusters alongside a complex microenvironment composed of cancer-associated fibroblasts (CAFs) and immune-infiltrated cells (Fig. 4 A, Fig. S7B, C). While cell type proportions were broadly comparable, the FFPE sample exhibited a higher fraction of tumor epithelial cells compared to the 10x Genomics sample (Fig. S7D). As before, the inferred phases accurately recapitulate known biological markers of cell cycle progression (Fig. S7E). Furthermore, the total transcript count per cell approximately doubles over the inferred cell cycle in both samples (Fig. S7F), and we observed a similar monotonic increase in cell area as a function of the inferred phase (Fig. S7G), providing orthogonal validations of the phase assignments.

**Fig. 4.**
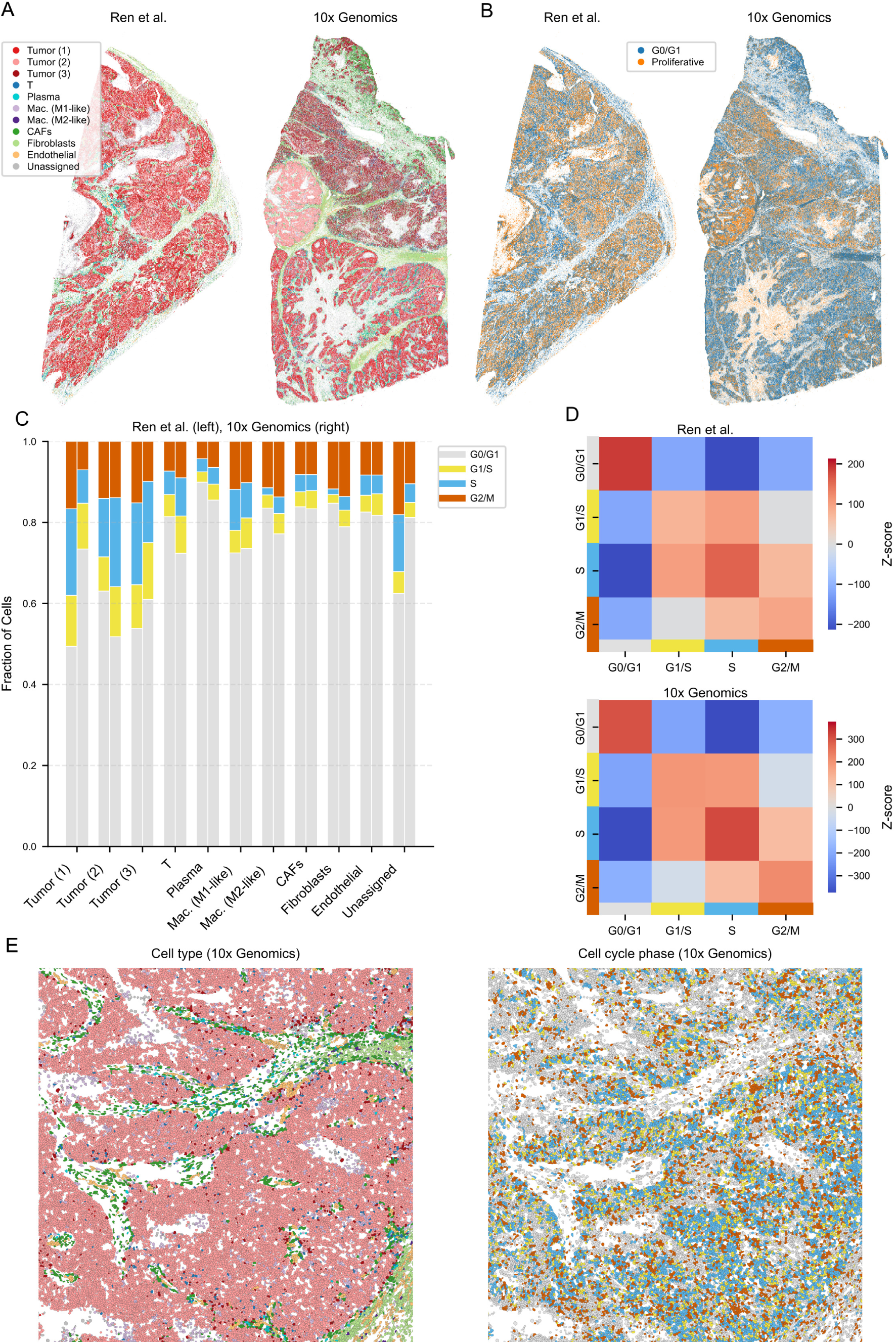
Spatial mapping identifies regions of phase coherence and proliferative hubs in ovarian cancer. Xenium Prime 5K In Situ Gene Expression for two human ovarian adenocarcinoma tissues: a metastatic high-grade serous carcinoma (FFPE) [37] and an adenocarcinoma sample (Fresh Frozen, 10x Genomics). **A.** Spatial cell-type maps showing 10x Genomics (top) and Ren et al. (bottom). Cell types were identified by Leiden clustering on the integrated *z*-space, distinguishing three tumor epithelial clusters, stromal, and immune infiltrated cells. **B.** Continuous cell-cycle phases inferred by CoPhaser projected onto the tissue coordinates. Cells are colored by their cycling status, with G1/S, S, and G2/M phases in orange and G0/G1 in blue. **C.** Fraction of cells in each cell cycle phase across the different cell types for 10x Genomics (left) and Ren et al. (right). **D.** Heatmaps of neighborhood *z*-scores displaying the spatial enrichment of cell cycle phases across the whole slide for Ren et al. (Top) and or 10x Genomics (bottom). **E.** High-resolution zoom of a highly proliferative Tumor (2) region showing cell type (left) and cell cycle phases (right).

By projecting these continuous phases back onto the tissue coordinates, we observed that proliferative cells (S/G2/M phases) are not stochastically distributed but instead occupy distinct proliferative regions within the tumor regions (Fig. 4B). These proliferative cells were significantly more abundant in malignant cell types compared to the stromal and immune cells, with sample-specific enrichment across the three epithelial sub-clusters (Fig. 4C).

To systematically characterize the spatial organization of these phases, we performed neighborhood enrichment analysis. In both samples, the matrices of neighborhood *z*-scores exhibited a prominent diagonal structure (Fig. 4D). This indicates that cells in a given cell cycle phase, particularly those in G1/S, S or G2/M, are significantly more likely to be surrounded by neighbors in the same or an adjacent cell-cycle phase. This local phase-clustering suggests a non-random spatial distribution where cell cycle progression is regionally coordinated, potentially reflecting localized signaling or clonal expansion. High-resolution inspection of the tissue architecture confirms these patterns, illustrating the clear spatial segregation between regions of active proliferation with patches of coordinated phases (Fig. 4E).

To ensure these patterns did not arise by chance, we compared our results to a stringent null model using random phase permutations of proliferative cells within each tumor cell type (Methods 4.9.6). This confirmed that the observed mean “patch size” of cells in the same phase is significantly higher than expected by chance across most tumor subtypes in both datasets (Fig. S7H, I), providing evidence of spatial synchronization in cell-cycle progression.

Together, these results demonstrate CoPhaser’s utility in uncovering phase-dependent spatial organization, identifying localized domains of cell-cycle synchronization that would be invisible to traditional single-marker assays.

### 2.7 CoPhaser identifies context-modulated circadian oscillations in the mouse aorta

We next evaluated the ability of our model to capture circadian rhythms in gene expression at the single-cell level, even though core circadian genes are lowly expressed, complicating the task. To this end, we analyzed a high-coverage scRNA-seq dataset from mouse aorta sampled every 6 h over a 24 h period [21], namely at external Zeitgeber (ZT) times ZT 0, 6, 12, 18. The performance of phase assignment in the circadian field is often assessed with the median absolute deviation (MAD; Methods 4.10.1) between predicted phase and external ZT. Here, the inferred phases closely tracked the ZT, with a MAD of 1.67 h across all cell types (Fig. 5A), with the lowest MAD for the smooth muscle cells (SMCs) and the highest for endothelial cells (Fig. 5B).

**Fig. 5.**
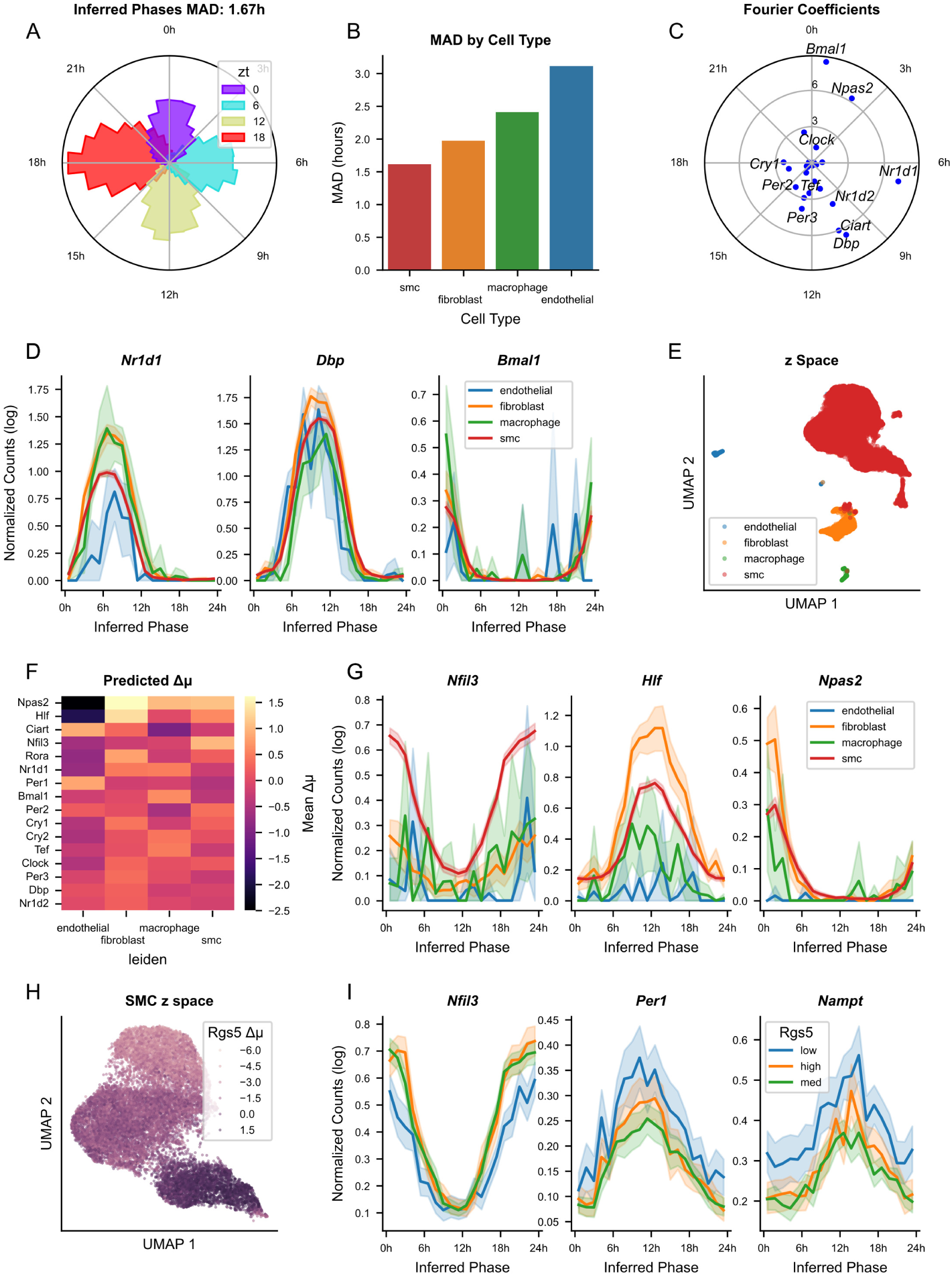
CoPhaser accurately predicts circadian phases and context-modulated gene expression profiles in the mouse aorta. Analysis of mouse aorta scRNA-seq data from [21] collected every 4h over a 24h light-dark cycle. **A.** Polar histogram of inferred phases for all cells, colored by Zeitgeber Time (ZT). The Median Absolute Deviation (MAD) between the predicted phase and the ZT is 1.67h. **B.** MAD between the inferred phase and the ZT for each cell type. **C.** Polar scatter plot of inferred Fourier coefficients for rhythmic input genes suggests a *bona fide* circadian clock, showing peak-to-trough amplitude (log) as radius and peak phase as angle. **D.** Reconstructed expression profiles of core clock genes (*Nr1d1*, *Dbp*, *Bmal1*) as a function of inferred phase, colored by cell type. Profiles were obtained by binning cells according to their inferred phase and averaging log-normalized expression (log(1+counts per 10,000)). **E.** UMAP projection of the inferred contextual *x*-space colored by cell type. **F.** Heatmap of inferred mean shifts (Δ*µ*) for rhythmic input genes across cell types. **G.** Reconstructed expression profiles of *Nfilh*, *Hlƒ*, and *Npas2* as a function of inferred phase, colored by cell type. Profiles computed as in **D**. **H.** UMAP projection of the inferred *x*-space colored by the inferred mean shift (Δ*µ*) of *Rgs5*, highlighting heterogeneity within smooth muscle cells (SMCs). **I.** Reconstructed expression profiles of *Nfilh*, *Per1*, and *Nampt* as a function of inferred phase, colored by SMC subtypes (Leiden clusters), corresponding to low, medium and high *Rgs5* expression levels. Profiles computed as in **D**.

The periodic gene expression trajectories (Fourier series) inferred by the model recapitulate the canonical circadian clock structure (Fig. 5C and [20]) and the inferred phases reveal rhythmic gene expression profiles across cell types (Fig. 5D). Notably, the overall MAD is lower than that obtained with *Tempo* [21], with particularly pronounced improvements for fibroblasts and macrophages (Table 1). These cell types are represented by relatively few cells but exhibit high amplitude clock gene expression rhythms (Fig. S8A). While minimizing MAD is not an explicit modeling objective, given that we expect genuine biological circadian phase heterogeneity across individual cells, achieving a lower MAD without using external time during training suggests that the context modeling in CoPhaser allows for a more accurate capture of the underlying clock structure. We attribute this improvement to the shared optimization across cell types, which stabilizes parameter estimation, especially for sparsely represented populations.

**Table 1.** Median Average Deviation (MAD, in hours) as a function of cell types. Mean over 5 independent CoPhaser runs, with the standard deviation indicated in parentheses. Our model was trained both with the same quality check filters as Tempo, and with an additional minimal number of 5000 UMIs per cell.

| Method | Gene set | SMCs | Fibroblasts | Endothelial cells | Macrophages |
| --- | --- | --- | --- | --- | --- |
| Tempo | All | 1.88 (0.03) | 2.53 (0.36) | 3.39 (0.4) | 2.99 (0.01) |
| Our Method | Core clock | 1.82 (0.03) | 2.30 (0.04) | 3.48 (0.29) | 2.50 (0.06) |
| Our Method<br>(high-quality cells) | Core clock | **1.73 (0.10)** | **2.01 (0.04)** | **2.84 (0.07)** | **2.29 (0.14)** |
| Random | - | 5.91 (0.02) | 5.79 (0.07) | 5.17 (0.28) | 5.34 (0.06) |

In parallel, the *z*-space faithfully captures cellular identity, separating the major vascular and immune cell types (Fig. 5E). Analyzing the inferred context-dependent shifts of the means reveals a cell-type-specific regulation of clock-related genes. In particular, *Npas2* and *Hlf* are predicted to be expressed at very low levels in endothelial cells (Fig. 5F). Inspection of reconstructed expression profiles confirms that both genes are nearly silent throughout the circadian cycle in endothelial cells (Fig. 5G), a pattern that is also evident at the pseudobulk level (Fig. S8B), while both genes display robust oscillations in other cell types. In contrast, *Nfil3* expression is largely restricted to smooth muscle cells (SMCs) (Fig. 5F,G). Analysis of the inferred amplitude scaling factor (*λ*) indicates that global circadian rhythmicity has the highest amplitude in smooth muscle cells and fibroblasts, while appearing slightly dampened in endothelial cells and macrophages (Fig. S8C).

A key benefit of CoPhaser is that it does not require prior clustering by cell type, but instead resolves cell heterogeneity directly from the data. Focusing on SMCs, the model resolves three distinct subpopulations characterized by differential expression of *Rgs5* (Fig. 5H, Fig. S8D). Using established marker genes [38], we identify the *Rgs5*-high and -intermediate clusters as originating from the descending thoracic aorta, whereas the *Rgs5* -low cluster corresponds to cells from the aortic arch (Fig. S8E). Beyond this anatomical stratification, the model predicts subtle, yet consistent differences in circadian gene regulation between these subpopulations, notably for *Nfil3*, *Per1* and *Nampt* (Fig. S8F). These differences are apparent both in the reconstructed single-cell expression profiles (Fig. 5I) and at the pseudobulk level (Fig. S8G), highlighting the model’s ability to resolve fine-grained context-dependent circadian regulation within closely related cell populations.

### 2.8 CoPhaser reveals menstrual cycle gene expression dynamics and endometriosis-associated alterations

We then applied CoPhaser to resolve the continuous dynamics of the human menstrual cycle. Throughout this cycle, endometrial stromal cells from the functionalis layer undergo extensive transcriptional remodeling under the influence of ovarian hormones [39], transitioning from proliferative endometrial stromal (eStromal) cells to decidualized stromal (dStromal) cells characteristic of the secretory phase. To infer a minimal cycling gene list, we used the mutual information *I* between the menstrual cycle phase and the gene expression (Method 4.8).

Consistent with this biology, the phases inferred closely match the annotated menstrual stages (Fig. 6A), with cells from the same subject assigned to highly similar phases (circular standard deviation 0.1*π*; Fig. S9A). The different stromal cell states, corresponding to distinct stages of the menstrual cycle, follow the expected temporal progression (Fig. 6B) and are well integrated in the *z*-space. This indicates that the dominant source of gene expression variation among stromal cells can be summarized by a single one-dimensional phase, with the notable exception of an additional variance component associated with the cell cycle, specifically observed in cells annotated as “eStromal proliferating” (Fig. 6C). The curated set of rhythmic input genes (Methods 4.8) spans all phases of the menstrual cycle; however, CoPhaser additionally identifies many rhythmic genes among the broader set of context genes (Fig. S9B). The reconstructed expression profiles reveal diverse temporal dynamics, ranging from sharp transitions—such as *CXCL14*, which is abruptly induced in mid–secretory phase, likely corresponding to the opening of the window of implantation [40], to smoother transitions, such as that observed for *MMP11* (Fig. 6E).

**Fig. 6.**
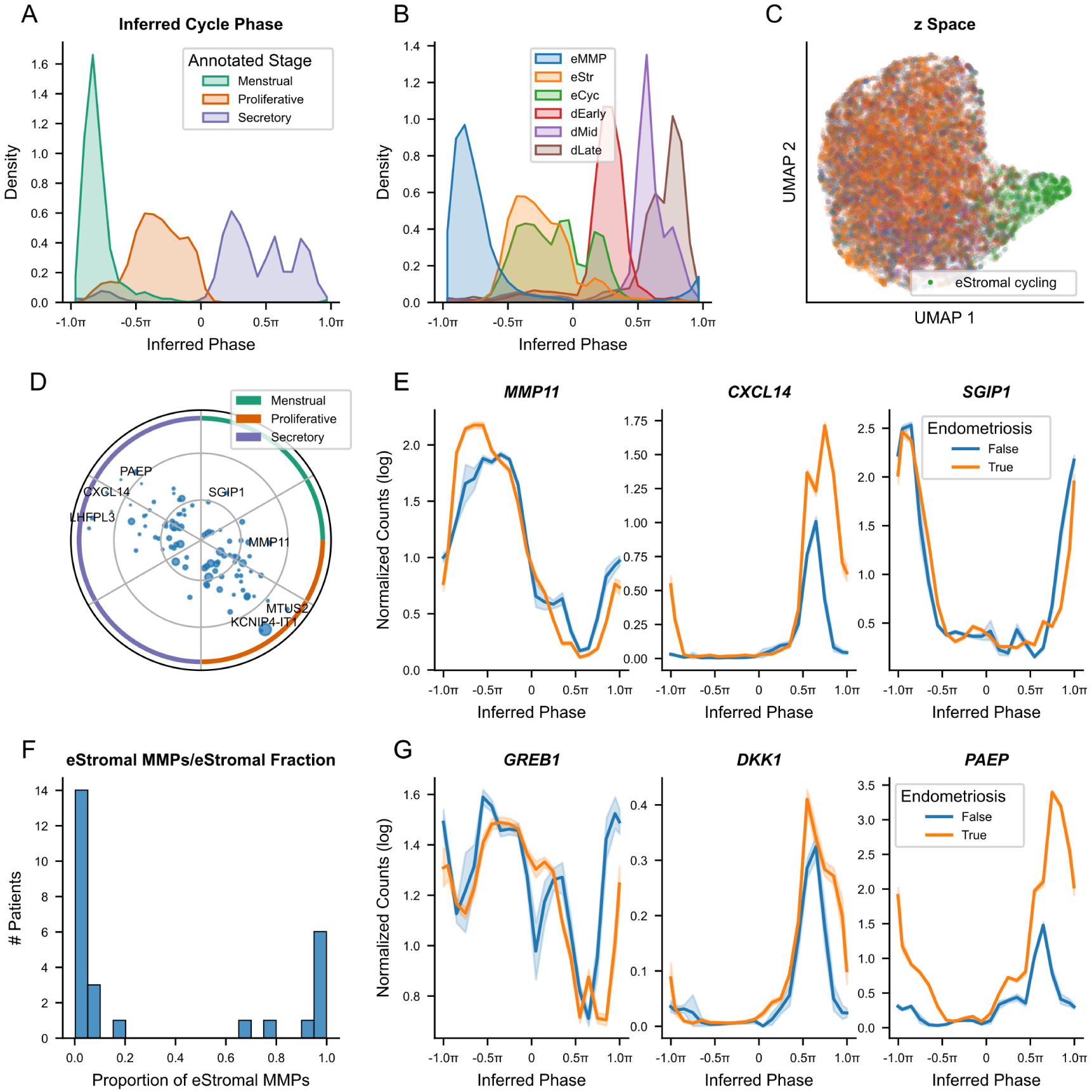
CoPhaser reveals menstrual cycle gene expression dynamics and endometriosis-associated alterations. Analysis of endometrium single-nucleus RNA-seq (snRNA-seq) data [39]. **A.** Distribution of inferred phases for stromal cells from donors without hormonal treatment, colored by the annotated menstrual cycle stage. **B.** Distribution of inferred phases for all stromal cells, colored by cell state. **C.** UMAP projection of the inferred contextual *x*-space, colored by cell state. **D.** Polar scatter plot of inferred amplitudes and phases for the set of rhythmic input genes, showing peak-to-trough amplitude (log) as radius and peak phase as angle. Point size is proportional to mean expression. Colored rings indicate the approximate menstrual cycle stages, corresponding to the color scheme of **A**. **E.** Reconstructed expression profiles of *MMP11*, *CXCL14*, and *SGIP1* as a function of inferred phase, comparing cells from healthy donors and donors with endometriosis. Profiles were obtained by binning cells according to their inferred phase and averaging log-normalized expression (log(1 +counts per 10,000)). **F.** Distribution of the proportion of eStromal MMPs cells (relative to the total eStromal population) across patients. **G.** Reconstructed expression profiles of *GREB1*, *DKK1*, and *PAEP* as a function of inferred phase, comparing cells from healthy donors and donors with endometriosis. Profiles computed as in **E**.

The original study described a distinct population of eStromal cells marked by high expression of matrix metalloproteinases (MMPs). CoPhaser predicts that this population is largely restricted to the menstrual phase, with minimal overlap with other eStromal cells (Fig. 6A). This prediction is further supported by the observation that few subjects harbor both populations simultaneously (Fig. 6F). We then examined differences between patients with endometriosis and control subjects. Because each individual spans only a narrow range of phases (Fig. S9B), and only a limited number of subjects cover each phase of the cycle (Fig. S9C), statistical power to detect deregulated genes across many subjects is limited. Among genes highlighted originally, *GREB1* and *DKK1* show modest differences limited to the late secretory phase. In contrast, analysis of inferred mean shifts revealed that *PAEP* is consistently more highly expressed in subjects with endometriosis compared to controls (Fig. 6G).

Finally, we leveraged the versatility of CoPhaser to analyze cell cycle dynamics occurring within the endometrium across the menstrual cycle. The model recovers the expected peaking phases and expression profiles of canonical cell cycle genes (Fig. S9D, E). The cell types identified as proliferative are consistent with the original annotations, with “Cycling” cells and “Cycling eStromal” cells exhibiting the highest proliferative activity (Fig. S9F). Notably, a subtype of endometrial perivascular cells (ePVs), annotated as “group 1b” based on *STC2* expression, also appears highly proliferative. As expected, the majority of proliferation occurs during the proliferative phase of the menstrual cycle (Fig. S9G). Across these proliferative cell types, we did not detect major differences in cell cycle gene expression between endometriosis patients and controls (Fig. S9H).

### 2.9 CoPhaser reveals coupling between the segmentation clock and the cell cycle

Finally, we applied CoPhaser to the segmentation (somite) clock, a biological oscillator that governs the periodic formation of somites during vertebrate embryonic development [41]. In the mouse embryo, somitogenesis proceeds with a periodicity of approximately 2h and is driven by oscillatory gene expression arising from three coupled signaling pathways: Notch, Wnt, and FGF, with *Hes7* acting as a core oscillator [42]. According to the classical clock-and-wavefront model, somite formation results from the interaction between this molecular clock and a posterior-to-anterior gradient of FGF and Wnt signaling: successive waves of *Hes7* expression propagate along the presomitic mesoderm (PSM) and trigger segmentation at the anterior boundary [41, 43]. To capture these dynamics, we analyzed a single-nucleus RNA-seq dataset of mouse embryos spanning roughly one embryo per somite stage up to the 34-somite stage. We selected PSM cells based on the original annotations [44] and used a curated list of mouse oscillatory genes [45] as rhythmic inputs.

The inferred *z*-space recapitulates the posterior–anterior axis of the PSM (Fig. 7A). A secondary source of variation correlates with embryo age, with younger embryos positioned to the right and older embryos to the left (Fig. 7B). This effect is dominated by a sharp transition after the 12-somite stage, which reflects a technical difference in sequencing depth, as embryos at early stages were drawn from a previous, deeper-sequenced study. Consistent with prior knowledge, *Hes7* and *Lfng* carry the largest information about the inferred somite clock phase (Fig. 7C) and display large-amplitude oscillations when plotted as a function of phase (Fig. 7D). Interestingly, although *Dkk1* is part of the Wnt pathway, expected to oscillate out of phase with *Hes7* in the posterior PSM and in phase in the anterior PSM, resulting in little net oscillation at the whole-PSM level [46], CoPhaser infers strong antiphasic oscillations at the mRNA level. To substantiate this result, we computed Pearson correlations between the raw log-normalized *Hes7* and *Dkk1* expression, which are negative in both posterior and anterior regions, consistent with the inferred dynamics. In contrast, *Axin2* is out of phase with *Hes7* in the posterior PSM but in phase in the anterior PSM (Fig. S10A). We further note that while *Per1* exhibits large oscillations, these are not reflecting a bona fide circadian clock, as neither *Bmal1* nor *Dbp* shows corresponding rhythmicity (Fig. S10B). Analysis of fitted oscillation amplitudes along the posterior–anterior axis reveals a progressive increase toward the anterior PSM, in agreement with previous observations[47]. Finally, it was even possible to reconstruct the traveling waves of gene expression across embryos. At the individual embryo scale, at each time point, a wave is traversing the PSM, leading to random phase offsets between the different embryos (Fig. S10D). Therefore, no coherent pattern is visible at the population level (Fig. 7F). However, after correcting for embryo-specific phase offsets (Fig. 7G), a clear posterior-to-anterior traveling wave emerges (Fig. 7H), covering approximately one full rotation over the PSM, as expected.

**Fig. 7.**
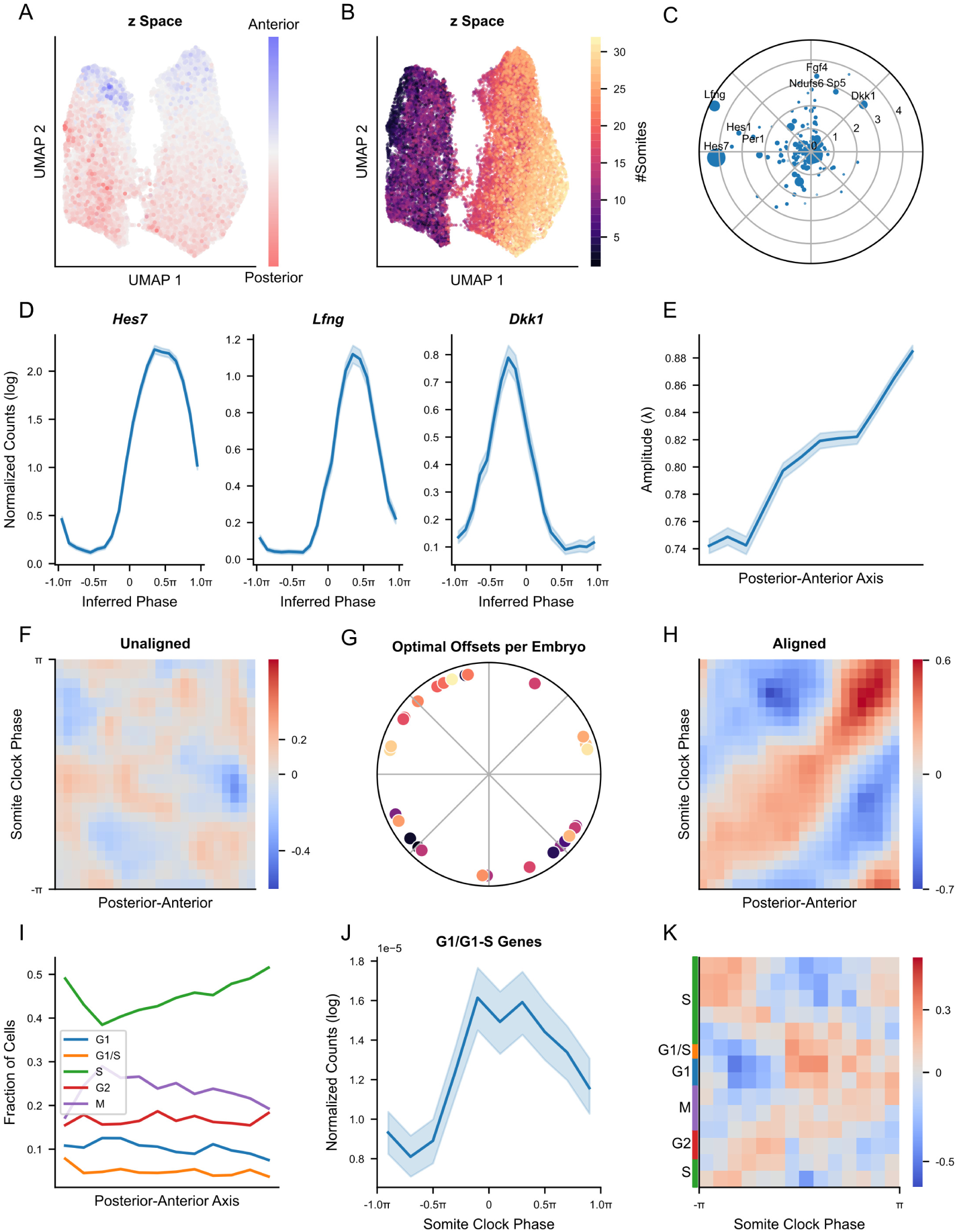
CoPhaser reveals coupling between the segmentation clock and the cell cycle. Single-cell analysis of mouse embryonic presomitic mesoderm (PSM) cells [44]. **A.** UMAP embedding of the inferred contextual *x*-space, colored by expression of *Wntha* (posterior marker) and *Meox1* (anterior marker). **B.** Same UMAP as in **A**, colored by the number of somites per embryo. **C.** Polar scatter plot of inferred amplitudes and phases for rhythmic input genes; point size scales with the product of oscillation amplitude and mean gene expression. **D.** Reconstructed expression profiles of *Hes7*, *Lƒng*, and *Dkk1* as a function of the inferred segmentation clock phase. Profiles were obtained by binning cells by phase and averaging log-normalized expression (log(1 + counts per 10,000)). **E.** Mean oscillation amplitude along the posterior–anterior axis. The axis is defined by quantile binning of the second UMAP dimension in **A**, performed independently for each embryo. **F.** Heatmap of pointwise mutual information (PMI) between somite clock phase and posterior–anterior position, computed using uncorrected phase estimates (i.e., without embryo-specific traveling wave correction). **G.** Polar plot of inferred embryo-specific traveling wave phase shifts, colored by somite number as in **B**. **H.** Same as **F**, after correcting somite clock phases using the embryo-specific phase shifts shown in **G**. **I.** Fraction of cells in each cell cycle phase as a function of posterior–anterior position. **J.** Median fraction of log-normalized counts contributed by genes peaking in the G1/G1–S phases, as annotated in Cyclebase [28]. **K.** Heatmap of pointwise mutual information between cell cycle phase and somite clock phase.

We next examined the relationship between the somite clock and the cell cycle. To do so, we used CoPhaser to infer a cell cycle phase in the same cells. The inferred cell cycle phase accurately recovers canonical cell cycle gene expression profiles (Fig. S10D), and the mean number of unique molecular identifiers (UMIs) per cell approximately doubles over the course of the inferred cycle (Fig. S10E). The PSM is characterized by a rapid cell cycle, with a short G1 phase and a proportionally extended S phase [48], which is reflected in the relative durations of the inferred cell cycle phases (Fig. S10F), similar to the other dataset of mouse embryonic cells (Fig. S4). In agreement with previous work [48], approximately 50% of PSM cells are assigned to the S phase. Along the posterior–anterior axis, we observe a decrease in cells at the G1/S transition from the posterior to the anterior PSM and a localized increase in M-phase cells accompanied by a reduction in S-phase cells at roughly one quarter of the axis.

Interestingly, genes peaking during G1 or at the G1/S transition display oscillatory expression with respect to the somite clock phase (Fig. 7J), peaking in phase with *Hes7*. Conversely, core somite clock genes (*Dkk1*, *Axin2*, and *Hes7*) also oscillate as a function of cell cycle phase, with elevated *Dkk1* expression from late S phase to mid G1 and a peak in *Hes7* at the G1/S transition (Fig. S10G). These reciprocal oscillations hint at a coupling between the cell cycle and the segmentation clock. This coupling is further supported by point-wise mutual information analysis (Fig. 7K), which reveals that the G1-to-S transition is least likely to occur when *Hes7* is at its minimum, and maximal probability with rising *Hes7* levels. Other cell cycle phases show weaker but phase-shifted associations, with almost no association during cell cycle phases with elevated *DKK1* expression.

## 3 Discussion

Biological cycles, including the cell cycle, circadian rhythms, the segmentation clock, and hormonal cycles, are intrinsic periodic processes operating within heterogeneous cellular environments. In vivo, these cycles are rarely perfectly synchronized across cells and are continuously modulated by cell identity, differentiation state, and disease context. As a result, cyclic gene expression programs cannot be faithfully represented as fixed trajectories shared across all cells. Instead, they are better described as periodic manifolds whose geometry is smoothly deformed by biological context, a structure we refer to as context-dependent periodic manifolds.

Most existing scRNA-seq-based cycle inference methods implicitly assume homogeneous dynamics across cells [14, 15, 22, 29, 49, 50]. While effective in relatively uniform systems, this assumption leads to confounding between latent phase and contextual variation in heterogeneous tissues, limiting interpretability and robustness. CoPhaser addresses this challenge by explicitly decomposing gene expression into a one-dimensional cyclic latent coordinate and a non-cyclic latent space that captures all remaining cellular variability. A mutual information-based constraint enforces statistical independence between these components, enabling a clean separation between periodic dynamics and context-dependent effects.

CoPhaser is broadly applicable across biological oscillators of distinct molecular origins and timescales, ranging from fast clocks such as the somite clock, cell cycle, and circadian rhythms to slower hormonal cycles. Beyond the assumption of periodicity, the model does not rely on cycle-specific constraints or predefined phase labels. Aside from an initial list of seed rhythmic genes, which may be incomplete and refined iteratively, the model learns both the phase definition and the deformation of the periodic manifold directly from the data.

This generality allows CoPhaser to unify the analysis of cyclic processes across contexts, while maintaining interpretability at the level of individual genes. By modeling both context-dependent shifts in baseline expression and context-dependent modulation of oscillation amplitude, the framework captures biologically meaningful deviations from an average cycle without compromising the coherence of the inferred phase.

Across cancer datasets, CoPhaser provides a principled way to distinguish constitutive transcriptional deregulation from phase-specific effects driven by altered proliferation dynamics, a distinction that is often obscured in bulk or pseudobulk analyses. In breast cancer, for example, we observed higher proliferation rates in triple-negative tumors compared to other subtypes, consistent with previous reports [51]. Importantly, CoPhaser allowed the disentanglement of this increased proliferative activity from genuine subtype-specific transcriptional programs.

This distinction is particularly relevant for interpreting apparent gene overexpression. Genes such as CDC20 appear upregulated in highly proliferative tumors, yet CoPhaser reveals that this effect arises primarily from phase-specific expression during S/G2/M rather than constitutive overexpression. For proteins with short half-lives, such as CDC20 [52], mRNA-level oscillations are expected to translate into protein dynamics, with important implications for therapeutic strategies. In particular, drugs targeting phase-restricted regulators may be less effective against slowly cycling persistent cancer cells [17].

Beyond individual genes, CoPhaser disentangled changes in cell-type abundance driven by differential proliferation from those driven by differential survival. In pediatric AML, we found that hematopoietic stem cells, while exhibiting low proliferation rates, can increase their relative abundance within the malignant compartment following treatment. This suggests enhanced treatment resistance rather than increased proliferation and provides a mechanistic explanation for the observed shift toward more primitive cell states at relapse [35]. More generally, these results highlight how explicit modeling of cell-cycle dynamics enables more accurate interpretation of compositional changes in cancer.

Applying CoPhaser to spatial transcriptomics in ovarian cancer revealed that proliferative cells are organized into synchronized cell-cycle phase patches. This “proliferative architecture” [18] suggests that cell-cycle progression is locally coordinated, likely reflecting the temporal integration of shared mitogen histories across cell generations [53]. These findings support the view that neighborhood composition is a primary driver of cell population dynamics [54]. The segregation of these domains from arrested populations likely involves conserved spatial CAF subtypes that orchestrate specific niches and immune exclusion [55]. Ultimately, these phase-stratified neighborhoods may define the boundaries of clonal expansions, where local microenvironments enforce synchronized proliferation decisions within the tumor ecosystem [56].

In the mouse aorta, CoPhaser recovered the circadian phase across cell types while simultaneously resolving cell-type-specific differences in baseline expression and oscillation amplitude of clock genes. The resulting context space exposes expected fine-grained differences in the regulation of the circadian programs [21, 22, 57] that would be difficult to detect with methods relying on either pseudobulk aggregation or cell-type-specific phase inference alone.

In the human menstrual cycle, CoPhaser captured transcriptional remodeling as a continuous trajectory, rather than a sequence of discrete states. The inferred phases support a view of hormonal cycling as a combination of smooth transitions punctuated by key transcriptional turning points that induce strong regulatory changes, similarly to what was described in [40]. This framework facilitates the discovery of disease-associated stromal gene expression modifications that are not readily accessible through pseudobulk analyses, particularly when individual subjects span only narrow temporal windows of the cycle. CoPhaser’s ability to generalize across cycles allowed to further characterize a subpopulation of stromal cells defined in [39] as highly proliferative.

In the developing mouse presomitic mesoderm, CoPhaser reconstructed oscillatory gene expression driven by the segmentation clock and enables the recovery of traveling waves underlying somitogenesis from snapshot data. Beyond phase inference, the model uncovers a coupling between the somite clock and the cell cycle, most prominently around the G1/S transition. These findings are consistent with recent imaging-based studies [48] and demonstrate that transcriptomic snapshots contain sufficient information to resolve interactions between multiple biological oscillators when analyzed with an appropriate structured model. Furthermore, inferring a phase from transcriptomic data allowed us to discover dynamics across hundreds of genes, removing the bias for the few selected reporter genes used in the imaging-based studies. However, we did not find differences between anterior and posterior coupling, which could be due to dynamical differences between the mRNA and the protein levels. Applying CoPhaser to a dataset specifically designed to analyze the somite clock, with more PSM cells and higher sequencing depth, could improve the resolution of the dynamics inferred.

One limitation of CoPhaser is its reliance on an initial set of rhythmic genes, although sensitivity to this choice is mitigated by the ability to infer rhythmicity among context genes. Detection of low-amplitude oscillations further depends on sequencing depth, particularly for sparse cell populations. While training costs exceed those of simpler approaches such as PCA-based phase inference, these remain tractable for most scRNA-seq datasets.

Model performance is sensitive to hyperparameter choices, which can be guided by biological prior knowledge and further stabilized through transfer learning across datasets. Finally, CoPhaser assumes the existence of a shared cycling gene program across contexts; extreme cases in which all cycling genes undergo strong, context-specific phase shifts, or where no shared rhythmic genes exist, would preclude a unified phase definition.

Overall, this work demonstrates the value of embedding biological topology directly into latent representations. By integrating explicit periodic structure with a flexible representation of cellular context, CoPhaser provides a general and interpretable framework for analyzing cyclic gene expression in complex biological systems. Its modular and open implementation enables reuse across datasets and biological domains, paving the way for comparative and integrative analyses of biological rhythms.

## 4 Methods

### 4.1 Model architecture

The model comprises two parallel encoders and two parallel decoders, designed to separate periodic dynamics from context-dependent variability. Two distinct gene sets are used as encoder inputs: a set of seed rhythmic genes (*X_f_*) informing the rhythmic encoder, and a broader set of highly variable genes (*X_z_*), which includes the rhythmic genes, informing the context encoder.

#### Rhythmic encoder

The rhythmic encoder processes the seed rhythmic genes *X_f_*. Raw counts are log-normalized as

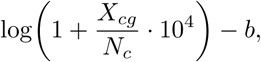

where *X_cg_*denotes the count of gene *g* in cell *c*, *N_c_* is the library size of cell *c*, and *b* is a baseline correction factor. The normalized expression is then projected through a three-layer fully connected neural network (Linear–Linear–Linear) into a two-dimensional latent space. Although functionally equivalent in expressivity to a single linear layer, this architecture has been shown to improve generalization when trained by gradient descent [58].

The resulting two-dimensional vector is normalized to unit length and used as the mean direction parameter *µ_φ_*(*X_f_*) of a Power Spherical posterior

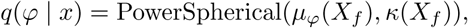

defined on the unit circle [25]. The concentration parameter *κ*(*X_f_*) is inferred via a separate neural network and informs on the uncertainty of the inferred phase.

#### Context encoder

The context encoder captures non-periodic sources of variation, such as cell type or technical effects, and closely follows a standard VAE architecture used in scVI [3]. Specifically, this encoder maps the variable genes *X_z_*to a latent variable *z* ∈ R*^dz^* using a multivariate normal posterior

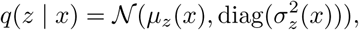

where *µ_z_*(*x*) and *σ_z_*(*x*) are produced by a two-layer fully connected neural network with GELU activations [59]. The dimensionality *d_z_*is a tunable hyperparameter; we typically use *d_z_* = 10 for complex biological contexts and *d_z_* = 2 for simpler settings.

#### Decoders and generative model

Three decoders are trained jointly from the latent spaces to reconstruct gene expression: (i) a rhythmic Fourier decoder capturing periodic expression patterns (*F_g_*), (ii) a context-dependent mean-shift decoder (Δ*µ*), and (iii) a context-dependent amplitude decoder (*λ*). The rhythmic component is decoded for all genes used by the model, not only the seed rhythmic genes, and is represented using a Fourier series with typically three harmonics (*N*_harm_ = 3) unless specified otherwise. The context-dependent decoders mirror the architecture of the context encoder.

We adopt a parametrized approach so that the expected gene expression is modeled as

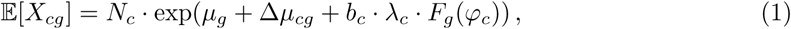

where *N_c_*is the library size, *µ_g_*is the gene-specific log-mean expression across the dataset, Δ*µ_cg_* is a context-dependent mean shift, and *λ_c_* is a context-dependent amplitude scaling factor derived from *z_c_*. The rhythmic component is given by

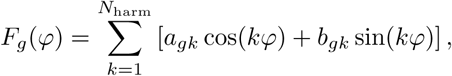

capturing the periodic expression pattern of gene *g*.

#### Context-dependent learning of corrections to the gene expression mean

To explicitly correct for context-dependent deformations of the periodic manifold, and for situations where the coverage of phases does not cover the cycle uniformly, we additionally decode a context-dependent shift in the mean 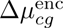 for the rhythmic encoder after an initial number of training epochs. This allows the model to adjust baseline expression levels and amplitudes prior to phase inference. The normalized counts used as input to the rhythmic encoder are then given by

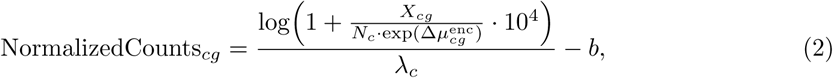

where 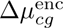 and *λ_c_* are both derived from the context latent variable *z_c_*.

### 4.2 Cycling classifier

For datasets containing substantial populations of non-cycling cells, such as adult tissues with G0-arrested cells, we include a cycling classifier to infer the probability that a cell is actively cycling. This classifier is implemented as a Relaxed Bernoulli (Gumbel–Softmax) encoder [60, 61], and is jointly trained with the rest of the model. Thus, while we illustrate the idea with G0 cells, this is generic and the nature of the G0-like state will depend on the cycle being considered. This encoder takes as input the gene expression *X* and the context latent variable *z*, and outputs a probability *b_c_*of being in the cycling state.

During training, samples are drawn from the Relaxed Bernoulli distribution to enable differentiable optimization. The temperature parameter is annealed exponentially over training to approximate a discrete Bernoulli distribution, allowing the model to make increasingly sharp cycling versus G0-like assignments.

When the prior probability of cycling is set to a value less than one, the expected expression is modeled as

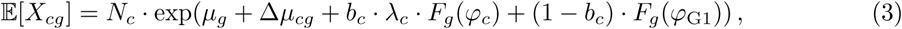

where *b_c_* ∈ {0, 1} denotes the cycling status of cell *c*, and *φ*_G1_ is a fixed reference phase corresponding to the G1 state when modeling the cell cycle (hence the notation), accounting for the transcriptional similarity between G0 and G1 cells. For other cycles, this reference can be chosen to represent arrested cycling, as for example, in a knock-out model of the circadian clock.

Although not strictly required, for datasets with a high fraction of non-cycling cells, we initialized the rhythmic decoder using Fourier coefficients inferred from the RPE1 dataset [29], and fixed *φ*_G1_ = −1.5 radians.

### 4.3 Mutual information neural estimation (MINE) network

To encourage the rhythmic phase *φ* and the contextual latent variable *z* to capture distinct sources of variation, we employ a mutual information neural estimator (MINE) [26]. The MINE network estimates the mutual information *I*(*φ*; *z*) by discriminating between samples drawn from the joint distribution *P* (*φ, z*) and samples drawn from the product of marginals *P* (*φ*)*P* (*z*), obtained by randomly permuting one variable within each minibatch.

The estimated mutual information is incorporated into the training objective as a penalty term, thereby discouraging statistical dependence between the phase and context representations. This regularization promotes disentangled latent spaces in which *z* captures non-periodic contextual variation while *φ* encodes cyclic dynamics.

In practice, the model is often able to capture contextual information more easily than a periodic structure. To prevent the context encoder from absorbing the cycle signal, we explicitly control gradient flow during MINE optimization. Specifically, when computing the mutual information loss, the projected periodic variable in *f*-space is detached from the computational graph, such that gradients from the MINE loss update the context encoder but not the rhythmic encoder. This asymmetric gradient application encourages *z* to relinquish cycle-related information while preserving the ability of the rhythmic encoder to capture the periodic manifold.

### 4.4 Loss function

The model is trained by maximizing the evidence lower bound (ELBO), augmented with additional regularization terms to promote biologically meaningful and stable solutions. The objective to be minimized is then

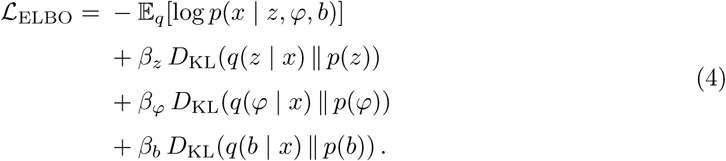

The reconstruction term log *p*(*x* | ·) is modeled using a negative binomial likelihood for count data. The Kullback–Leibler (KL) divergence terms regularize the approximate posteriors toward their respective priors: *p*(*z*) is a standard multivariate normal distribution *N* (0*, I*), *p*(*φ*) is a uniform distribution on the circle, and *p*(*b*) is a Bernoulli prior representing cycling status. The weighting coefficients *β_z_*, *β_φ_*, and *β_b_* implement a *β*-VAE framework [62]. In practice, *β_φ_* was set to 0.1 to prevent premature collapse of the phase posterior to a uniform distribution, which would hinder learning of cyclic structure, while *β_b_*was set to 10 to encourage the exclusion of non-cycling (G0-like) cells, which are transcriptionally similar to G1-like cells.

To further disentangle periodic dynamics from contextual variation, we include a mutual information penalty between the phase variable *φ* and the contextual latent variable *z*, for cells classified as cycling:

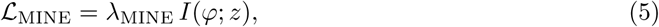

where *I*(*φ*; *z*) is estimated using a mutual information neural estimator (MINE), and *λ*_MINE_ controls the strength of the penalty. This term encourages *φ* to capture periodic structure while *z* encodes non-periodic contextual effects. Complete independence is not enforced, as some biological contexts (for example, cell type) are intrinsically associated with cycling propensity.

In the absence of additional constraints, the model can converge to degenerate solutions in which inferred phases collapse to a narrow range. To prevent this behavior, we introduce a circular entropy regularization term that encourages a uniform phase distribution early during the training. The phase distribution is estimated using a differentiable circular histogram with *N*_bins_ bins. Let *c_k_* denote the center of bin *k* and *w* = 2*π/N*_bins_ the bin width. For a batch of phases 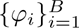, the contribution of phase *φ_i_* to bin *k* is computed using a Gaussian kernel on the circular distance:

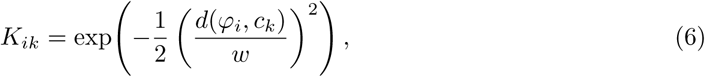

where *d*(*φ_i_, c_k_*) denotes the shortest angular distance between *φ_i_* and *c_k_*. The normalized probability mass of bin *k* is

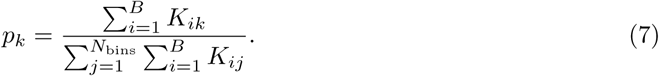

The circular entropy is then defined as 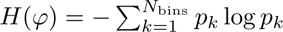, yielding the loss term

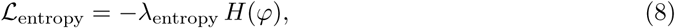

where *λ*_entropy_ controls the strength of the regularization. Importantly, to avoid over-constraining the phase distribution, this term is exponentially annealed during training (Methods 4.5). Its purpose is to route the optimization towards biologically relevant solutions early during the training.

Finally, to stabilize the geometry of the periodic latent space, we constrain the two-dimensional output *u* of the rhythmic encoder (prior to normalization onto the unit circle) to lie close to the unit circle. This is achieved by penalizing both the mean squared deviation of the radius from unity and its variance:

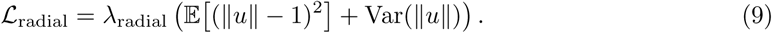

### 4.5 Training procedure

The model is trained for 200 epochs using the Adam optimizer with a learning rate of 0.01. To promote stable convergence, we adopt a staged training strategy. As described above, the weight of the circular entropy loss decays exponentially according to

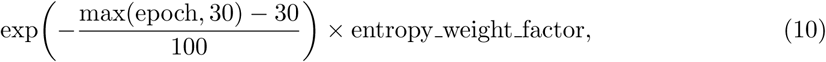

where entropy weight factor is a tunable hyperparameter.

To further mitigate training instabilities, corrections for the mean and amplitude in the rhythmic encoder 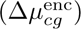 are enabled only after epoch 50, and the rhythmic amplitude parameter (*λ*) is optimized only from epoch 100 onward. For the cycling classifier, the temperature of the Relaxed Bernoulli distribution is annealed from a high initial value to a low final value over the course of training.

The mutual information neural estimator (MINE) is trained in parallel with the variational autoencoder (VAE), also using the Adam optimizer, to continuously estimate and minimize the mutual information between the dynamically evolving *z* - and *f*-spaces.

Although the model can be trained from random initialization, and was trained as such for cell-cycle datasets with low fraction of G0 cells and for menstrual cycle data, we found that non-random initialization of the Fourier coefficients was beneficial for more challenging settings. These include contexts with a low fraction of cycling cells and circadian datasets, in which only a small number of lowly expressed genes exhibit oscillatory behavior. In these cases, prior initialization was not strictly required but substantially reduced the likelihood of convergence to biologically implausible solutions, such as aberrant inferred peaking phases or flat histone expression profiles.

Initialization of Fourier coefficients when specified were taken from the RPE1 dataset from [29] for cell-cycle analyses, and from [20] for circadian analyses, using only the genes shown in Fig. 1B. The Fourier coefficients were held fixed during the initial training epochs and subsequently unfrozen to allow adaptation to the target dataset.

### 4.6 Phase definition

Because phase inference is invariant to global phase rotations and phase orientation, an external convention is required to fix both the origin and the direction of the inferred phase.

For the cell cycle, the model automatically determines the phase orientation by enforcing the expected ordering of peak expression of canonical cell-cycle genes, such that genes peaking in G1 precede those peaking in S phase, and so on. We fixed the origin in this work such that −*π* and *π* correspond to the M phase. This alignment was achieved using the mean UMI count per cell, which exhibits a characteristic decrease during mitosis.

For the circadian cycle, the indeterminacy was resolved using the external sampling time, and for the menstrual cycle, we fixed the phase using the annotated menstrual stages.

### 4.7 Simulations

To simulate biologically plausible gene expression profiles for the cell cycle, we used the Fourier coefficients of the rhythmic gene set fitted on the two batches of RPE1 scRNA-seq from [29]. We then drew a phase and context variable ranging from 0 to 1, from uniform distributions for each of the 10,000 cells. For one third of the rhythmic genes, we drew a maximal mean shift from a uniform distribution ranging from −5 to 5 (log), and shifted the mean gene expression of each cell based on the formula

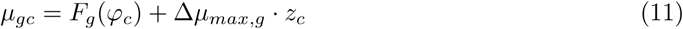

with *µ_gc_* the expected log fraction of count of gene *g* for the cell *c*, *F_g_*(*φ_c_*) the Fourier coefficient of the gene *g* evaluated at the phase *φ_c_*, Δ*µ_max,g_* the maximal mean shift for the gene *g* and *z_c_* the context variable for the cell *c*.

Furthermore, we scaled the Fourier coefficient by an amplitude factor ranging from 0.8 to 1.7 linearly based on the context value. We then simulated 2000 context variables, with randomly drawn mean log fraction of counts, and maximal fold changes randomly drawn between −3 and 3, which modifies the expected log fraction by scaling them linearly between their maximal fold change and their basal expression, according to the context variable. We then transformed these expected log fractions into counts using a negative binomial noise with as mean the expected fractions multiplied by a target of 10,000 counts per cell, and a dispersion parameter of 0.1.

### 4.8 Gene sets selection

The model requires two gene sets: a rhythmic gene set, known to oscillate with the cycle of interest, and a context gene set, providing a context sufficient to recapitulate the different populations of cells found in the dataset. For the context genes, we always used the 2000 most variable genes in the dataset using the function *highly variable genes* from *Scanpy* with the flavor *Seurat* [63]. For the rhythmic gene set we tailored the list for each of the cycles: for the cell cycle, we use a list of 98 cell cycle genes from [29], for the circadian cycle, we used a list of 25 core clock genes created by [21], for the somite clock we used a list of 175 genes created by [45], and for the menstrual cycle, we used a list of 93 genes, obtained by selecting the 3000 most variable genes in the stromal cells, and taking the genes with an *I* greater than 0.1 between their counts and the menstrual cycle state using the *Sklearn* function *mutual info classif* [64]. All the rhythmic gene lists can be found in Supplementary Table 1 (excel file).

#### 4.8.1 Rhythmic gene selection from fitted context genes

To select the genes added to the rhythmic gene set using the two-gene fit (See Results 2.3), we sorted the context genes according to their mean fraction of counts multiplied by the exponential of the fitted peak-to-peak amplitude.

### 4.9 CoPhaser inference on multiple datasets

#### 4.9.1 RPE1 cells from Battich *et al*

This dataset of RPE1 cells was sequenced using single-cell EU-labeled RNA sequencing, which is based on CEL-Seq2 [8]. We used the original mapping (Table 2) and removed cells with less than 5% of unspliced counts, more than 70% of unspliced counts, and less than 2,000 UMIs per cell. 2,305 cells passed the quality check (QC) cut-offs and were kept for the training of our model, which was performed from scratch.

**Table 2.** List of datasets used in this study and their corresponding access locations.

| Name | Cycle | Link | Original Paper |
| --- | --- | --- | --- |
| RPE1 | Cell cycle | GSE128365 | [8] |
| RPE1 | Cell cycle | GSE250148 | [29] |
| Mouse embryo | Cell cycle | GSE176588 | [16] |
| Breast cancer | Cell cycle | GSE161529 | [70] |
| Pediatric AML | Cell cycle | GSE235063 | [35] |
| Ovarian cancer (spatial) | Cell cycle | 10x Genomics | (2026, January) |
| Ovarian cancer (spatial) | Cell cycle | Their website | [37] |
| Aorta | Circadian cycle | dropbox | [21] |
| Endometrium | Menstrual cycle | E-MTAB-14039 | [39] |
| Mouse embryo | Somite clock | Their website | [44] |

#### 4.9.2 RPE1 cells from Lederer *et al*

For this dataset [29], we mapped raw sequencing reads (Table 2) to the reference genome using *Cell Ranger* [65] (10x Genomics). Spliced and unspliced transcript counts were then extracted and demultiplexed using the *Velocyto* pipeline [66]. We then removed cells with less than 10% of unspliced counts, more than 42% of unspliced counts, fewer than 4,000, and more than 40,000 UMIs per cell. 15,237 cells passed the QC cut-offs and were kept for the training of the model, which was performed from scratch.

#### 4.9.3 Mouse embryonic cells from Salmen *et al*

For this dataset [16], raw sequencing reads (Table 2) were mapped to the reference genome using *Cell Ranger* [65] (10x Genomics). Spliced and unspliced transcript counts were then extracted and demultiplexed using the *Velocyto* pipeline [66]. We then removed cells with less than 30% of unspliced counts, more than 70% of unspliced counts, fewer than 1,500, and more than 50,000 UMIs per cell. 38,913 cells passed the quality check (QC) cut-offs and were kept for the training of the different models. We trained our model from scratch.

For Seurat’s discrete cell cycle phase assignment, we used its original implementation in R, and the list of genes from [67], as advised in their tutorial [68]. To assess the performance of DeepCycle [15], we tried to fit the model using the filtering method implemented using the intersection between the 2000 most variable genes and our list of cycling genes. However, the results were worse than directly using our 98 cycling gene list without filtering, reason why we used the latter. We inferred cell cycle phases using Tricycle [50], using its original implementation in R, analyzing all cell types individually, using the original author’s annotation. For Cyclum [14], we let the model find the best number of linear dimensions, which it found to be one, and used the hyperparameters that they used for mouse embryonic stem cells. For the PCA-derived phase inference, we first selected the genes in our cell cycle gene set and normalized the gene counts using Scannpy *normalize total* function, with 10,000 as the target, and took the log of 1 plus the normalized counts. Then we mean-centered each gene’s log-normalized counts, either estimating the mean using the full dataset or using the original authors’ cell types annotation. We then selected the first two PCs to infer the cell cycle phase. Using the original authors’ S-phase annotation, we also selected the first and third PCs, which appeared to contain more variance associated with the cell cycle. We tested scPrisma [69], but not all cells could fit in the GPU used and using only 20% of the dataset led to inconclusive results. Furthermore, scPrisma fits can be improved using prior knowledge, but Seurat’s estimation of the phase does not constitute a good prior in this case. We compared all these methods with a random phase assignment, for which the cell phases are drawn from a uniform distribution.

#### 4.9.4 Breast cancer atlas from Pal *et al*

We analyzed this breast cancer atlas [70], using the original mapping and cell type annotation (Table 2). We dropped the sample ER 0001 because the cell barcode from the count matrix do not match the one containing the cell type annotation. We merged the data from the three tumor subtypes and selected the tumoral epithelial cells based on the original annotation. We used the cells that passed the original QC, leading to 85,542 cells. We trained the model starting with the Fourier coefficients obtained from the dataset of RPE1 cells from Lederer *et al.*, frozen for the first 20 epochs.

#### 4.9.5 Dataset of pediatric AML cells from Lambo *et al*

We analyzed this longitudinal dataset of pediatric AML [35], using the original mapping and cell type annotation (Table 2). We merged all the patients and timepoints and selected the normal cells that had more than 2,000 UMIs and less than 6,000 UMIs and Malignant cells that had between 6,000 and 20,000 UMIs. To reduce training time, we kept only 50% of these cells, leading to 90,891 cells and a training time of less than 15 minutes. Similarly as before, we started the training of the model with the Fourier coefficients obtained from the dataset of RPE1 cells from Lederer *et al.*, frozen for the first 20 epochs.

#### 4.9.6 Spatial transcriptomics data from 10x Genomics and Ren *et al*

Spatial transcriptomics data (Xenium 5k) were obtained from 10x Genomics (10xG) website and from [37] (REN) (Table 2). These datasets were chosen because of their high sequencing depth (median transcripts ¿700) compared to other spatial datasets. The 10x Genomics Xenium output bundle was converted to .zarr format using the spatialdata framework, while the Ren *et al.* dataset, comprising raw transcripts (parquet), count matrices (h5ad), and segmentation masksm, was assembled into a SpatialData object and saved as .zarr. l Post-filtering, REN contained 372’113 cells (median transcripts per cell: 747 transcripts), and the 10xG contained 1’062’506 cells (median transcripts per cell: 1’467). For joint modeling, count matrices were merged and processed using CoPhaser with the cycling classifier. The Fourier decoder was initialized with the Fourier coefficients obtained from the dataset of RPE1 cells from Lederer *et al.*, which remained frozen for the initial 20 epochs; the 2’000 most variable genes were utilized for context. Harmony [71] was applied to the resulting latent *z*-space to obtain a batch-corrected latent context. A *k*-nearest neighbor (*k*-NN) graph was then constructed on the Harmony embeddings, followed by clustering via the Leiden algorithm (resolution: 0.5). Cell types were annotated by identifying differentially expressed genes and cross-referencing them with canonical markers from the CellMarker 2.0 database [72]. To investigate spatial organization, spatial neighbor graphs were constructed using Delaunay trian-gulation via squidpy.gr.spatial neighbors at the whole-slide level. Neighborhood enrichment analysis was performed to calculate *Z*-scores for spatial co-occurrence between cell-cycle phases using squidpy.gr.nhood enrichment. We further applied a custom permutation-based analysis to quantify the spatial clustering of phases of proliferative cells. Within cell-type-specific spatial clusters containing ≥ 100 cells, we identified spatially connected components (termed phase patches) for each cell cycle phase (G1/S, S, G2/M) using the spatial adjacency graph. For each cluster and phase, the mean patch size (average number of cells per connected component) was computed. To generate a null expectation, we performed permutations where cell-cycle phase labels were randomly shuffled exclusively among cycling cells (G1/S, S, G2/M), and within each cell-type cluster independently to avoid cell-cycle phase frequency biases, while preserving their spatial positions. Statistical significance was assessed using paired *t*-tests comparing observed versus expected mean patch sizes across all clusters within each tumor type.

#### 4.9.7 Dataset collected around the clock from Auerbach *et al*

We analyzed the dataset collected around the clock from the mouse aorta [21], because its high quality (median UMI around 10,000 counts) allows circadian phase inference at the single cell level. The cell types annotation and counts were downloaded directly from the original work (Table 2). Based on the histogram of the number of counts per cell, we added a second filtering on their data by removing the cells with fewer than 5,000 UMIs. For comparison with their obtained median absolute deviation (MAD), we used both the subfiltered dataset and the original one and reported the results on both (Table 1). We used the same clock-related gene sets as the original authors, which can be found in Supplementary Table 1.

#### 4.9.8 Dataset from the human endometrium from Mareckova *et al*

We analyzed the dataset of single-nucleus RNA-seq from human endometrium by [39]. The authors sequenced cells from snap-frozen samples of superficial endometrial biopsies from 63 donors, with and without endometriosis. We used the mapping and counts provided by the authors available at ArrayExpress (Table 2), and selected the stromal cells for the analysis of the menstrual cycle, and randomly selected one quarter of the dataset, leading to 30,536 cells. The cycling gene list was obtained as explained above (4.8). For the cell cycle, we randomly sampled one-third of the dataset for the cell cycle analysis, leading to 93,674 cells.

#### 4.9.9 Dataset of mouse embryos from Qiu *et al*

We analyzed the dataset of single-nucleus RNA-seq from mouse embryos by [44]. The authors sequenced 12.4 million nuclei from 83 embryos, with the 32 first embryos containing 0 to 34 somites. We used the mapping and counts provided by the authors (Table 2), and their annotation. We selected the cells annotated as *Tbx6* positive, integrated them using harmony [71], and reselected more precisely the cycling cells, based on clusters positive for *Tbx6*, *Hes7*,*Dll1*, and *Dkk1*, leading to 25,130 PSM cells. The cycling gene list was obtained from [45], and can be found in Supplementary Table 1. The model was trained using the phase provided by [45] as initialization of the first harmonic of Fourier coefficients, with an approximate amplitude fitted on the data from the original authors.

### 4.10 Evaluation metrics

#### 4.10.1 Median absolute deviation (MAD)

To evaluate model performance on the mouse aorta dataset collected around the clock, we followed the original study and quantified accuracy using the median absolute deviation (MAD) between the inferred phase and the external sampling time. Specifically, MAD is defined as

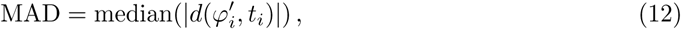

where 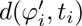 denotes the circular distance between the inferred phase 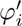 and the external time *t_i_*of cell *i*.

Because both the origin and direction of the inferred phase are arbitrary, we computed MAD after optimally aligning the inferred phases to external time by selecting the shift and orientation that minimized the deviation, independently for each cell type.

As noted previously, a low MAD is not always expected, as it would imply perfect synchrony of circadian phases across individual cells. Nevertheless, achieving a lower MAD without incorporating external time into model selection or training provides evidence that the inferred phases capture biologically meaningful circadian structure.

#### 4.10.2 Jensen–Shannon divergence

To assess agreement between inferred phases and the binary S-phase annotation provided by the original authors [16], we computed the Jensen–Shannon (JS) divergence between the phase distributions of S-phase and non–S-phase cells. The JS divergence is a symmetric and smoothed variant of the Kullback–Leibler divergence, defined as

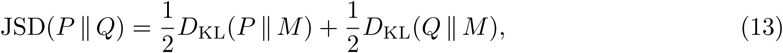

where 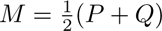 is the mixture distribution.

To obtain a proper distance metric, we report the square root of the JS divergence, which satisfies the triangle inequality [73]. Using a base-2 logarithm, the resulting Jensen–Shannon distance is bounded between 0 and 1, with 0 indicating identical distributions and 1 indicating completely non-overlapping distributions.

### 4.11 Use of Large Language Models

Some code snippets used in this work were generated using LLMs, reviewed, and modified by the authors. Furthermore, the authors used LLMs to improve the clarity, coherence, and linguistic quality of the text. After using this tool, the authors reviewed and edited the content as needed and take full responsibility for the content of the publication.

## 5 Supplementary Material

### 5.1 Supplementary Table S1

Excel file containing the rhythmic gene lists for the four cycles.

### 5.2 Code Availability

CoPhaser is implemented in Python and freely accessible at https://github.com/gityves/CoPhaser. Source code, installation instructions, and notebooks to replicate the figures are available in the repository. Details regarding the the analyses in all the figures, such as the model hyperparameters, can be found in the notebooks.

## Acknowledgments

We thank members of the Naef lab for insightful discussions. This work was supported by the Swiss National Science Foundation (individual grant 310030B 201267 to F.N.) and by the Chan Zuckerberg Initiative (CZI) Single-Cell Biology Data Insights program (DI3) grant “Space-time Regulation of Biological Cycles in Cancer” awarded to Felix Naef and Nacho Molina.

## Supplementary Figures

**Fig. S1.**
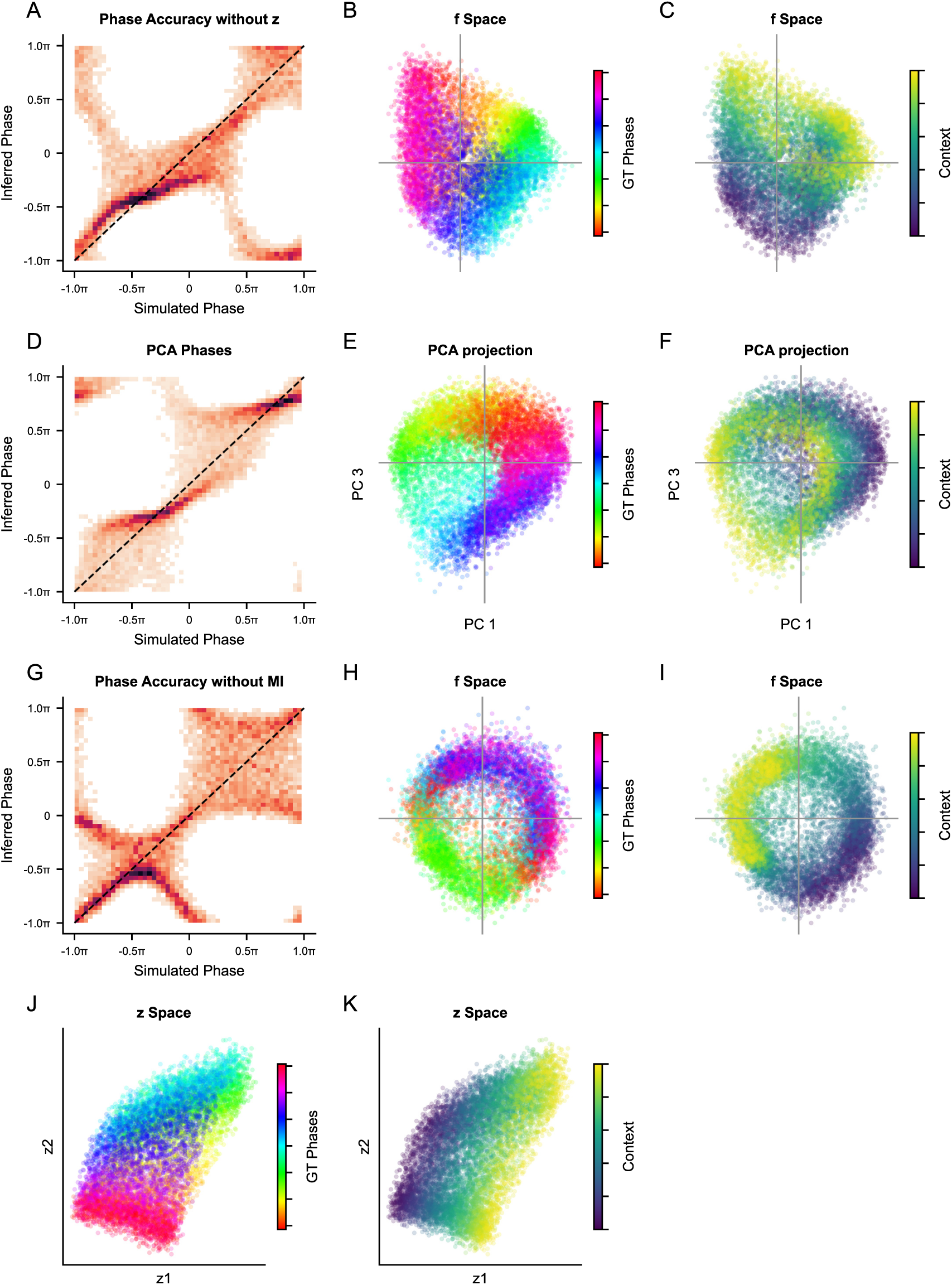
Ablation study validates the dual latent space architecture and independence constraints. **A–C.** Results on simulated data when the contextual *x*-space is removed from the model. **A.** Comparison between inferred phases and the simulated ground truth (GT), colored by local density. **B, C.** Projected *ƒ*-space colored by the GT phase (**B**) and by the one-dimensional contextual variable (**B**). The axis intersection indicates the (0,0) center. **D–F.** Principal component analysis (PCA) of the simulated data, projected onto the two components capturing most of the cyclic variance (PC1 and PC3). Panels are colored by local density (**D**), GT phases (**E**) and contextual variable (**F**). **G–K.** Results on simulated data without the Mutual Information Neural Estimation (MINE) to enforce independence between latent spaces. **G.** Comparison between inferred phases and GT phases, colored by local density. **H, I.** Projected f-space colored by GT phase (**H**) and contextual variable (**I**). **J, K.** z-space colored by GT phase (**J**) and contextual variable (**K**).

**Fig. S2.**
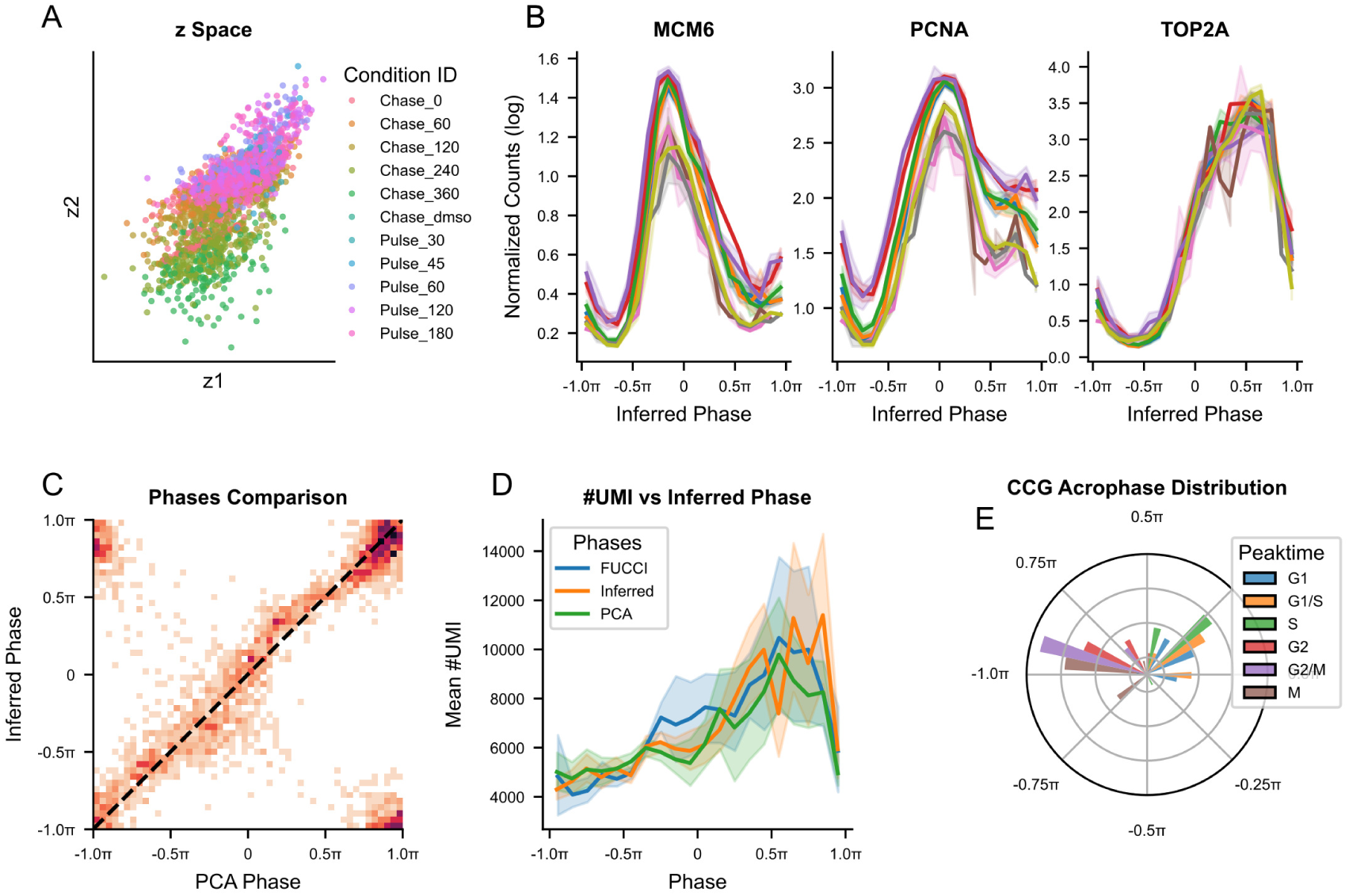
Validation of context-corrected inferred cell-cycle phases. Extended analysis of the RPE1 scEU-seq dataset [8]. **A.** Inferred contextual *x*-space, colored by experimental condition (pulse/chase durations). **B** Reconstructed gene expression profiles for selected cell-cycle genes (*MCM6*, *PCNA*, *TOP2A*) as a function of inferred phase, colored by condition. Profiles were obtained by binning cells according to their inferred phase and averaging log-normalized expression (log(1 + counts per 10,000)). **C** Comparison between phases inferred by CoPhaser and phases obtained from principal component analysis (PCA), colored by local density. **D** Mean number of unique molecular identifiers (UMIs) per cell as a function of phase for FUCCI-derived, model-inferred, and PCA-derived phases in the chase condition. **E** Polar histogram of inferred peak phases for cell-cycle genes, colored by their annotated peak phase according to Cyclebase [28].

**Fig. S3.**
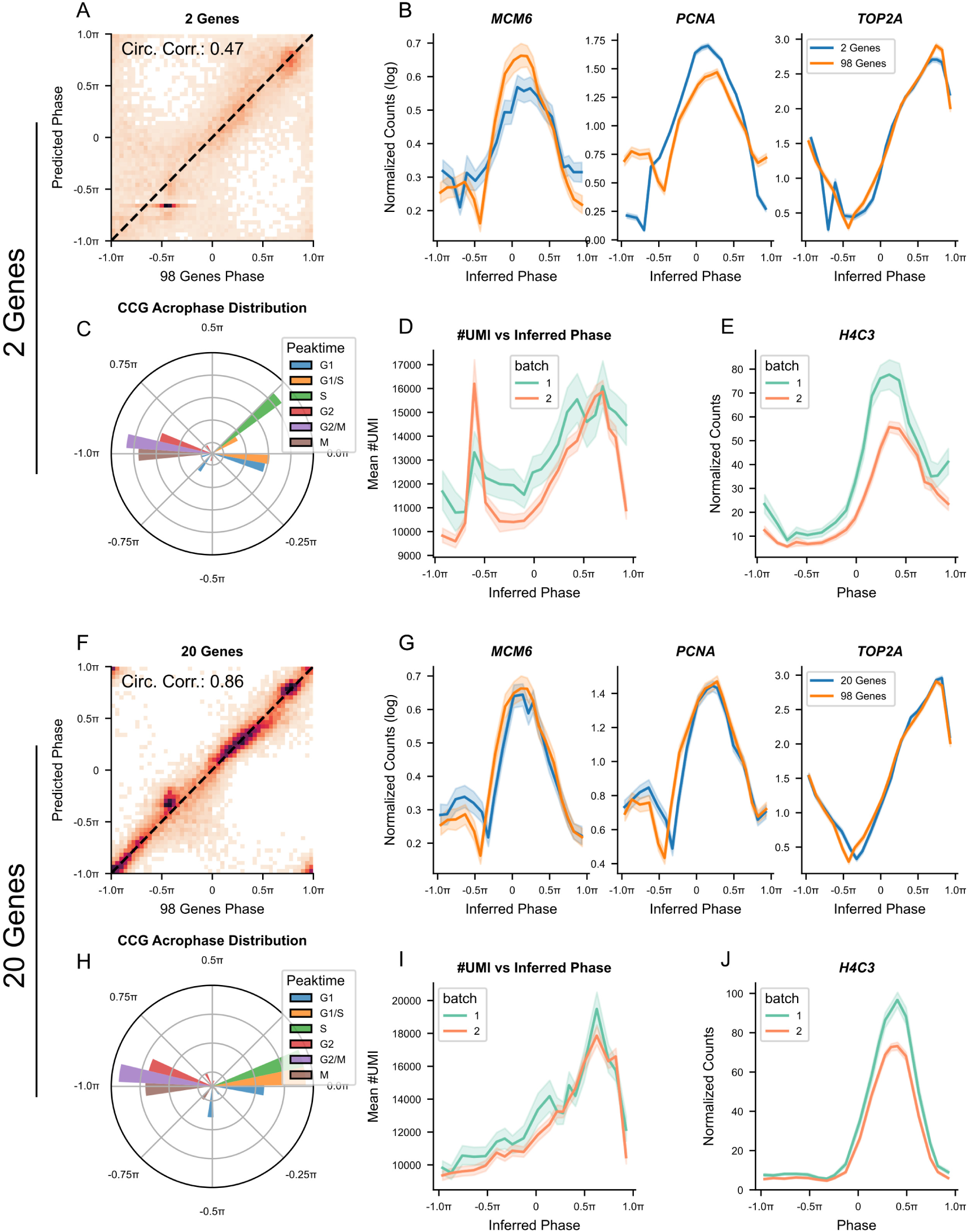
Iterative refinement of the cycling gene set enables robust phase inference. Extended analysis of the RPE1 scRNA-seq dataset [29]. **A–E.** Initial model performance when fitted using only two rhythmic input genes (*TOP2A* and *PCNA*). **A.** Comparison between phases inferred by the 2-gene model and phases inferred by the full 98-gene model, colored by local density. Circular correlation = 0.47. **B.** Reconstructed expression profiles of selected cell-cycle genes (*MCM6*, *PCNA*, *TOP2A*) as a function of phase, comparing the 2-gene model and the full 98-gene model. Profiles were obtained by binning cells according to the inferred phase and averaging log-normalized count log(1 + counts per 10,000). **C.** Polar histogram of inferred peak phases for cell-cycle genes, colored by their annotated peak phase according to Cyclebase [28]. **D.** Mean number of unique molecular identifiers (UMIs) per cell as a function of the inferred phase, colored by batch. **E.** Mean fraction of counts computed from histone genes as a function of phase inferred, colored by batch. **F–J.** Improved model performance using a refined set of 20 genes. Genes were selected based on a score combining mean expression and inferred amplitude (see Methods). Panels are analogous to **A–E** and show the removal of artifacts observed in the initial model.

**Fig. S4.**
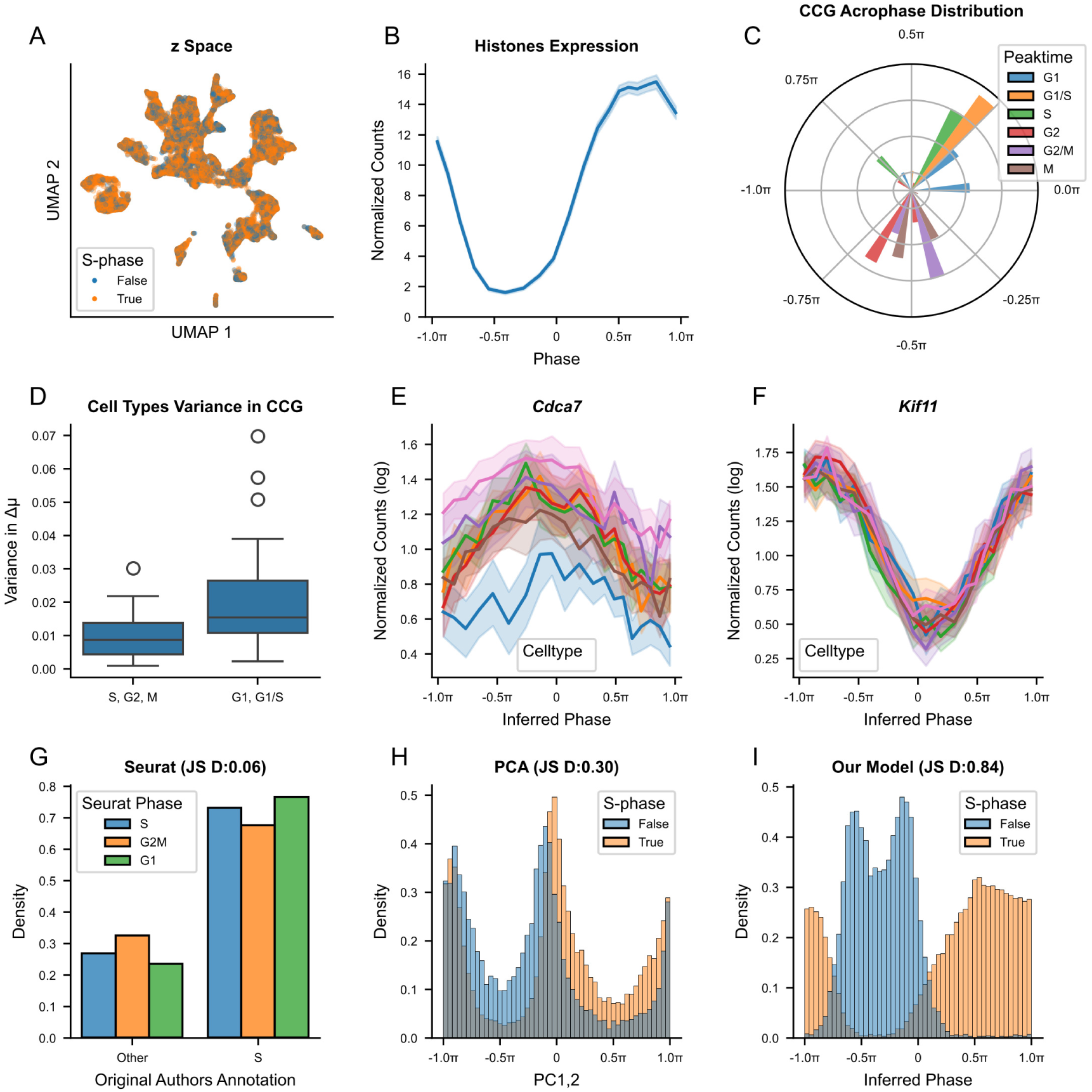
Validation of context-corrected cell-cycle inference in mouse embryonic cells. Extended analysis of the mouse embryonic VASA-seq from [16]. **A** Projected contextual *x*-space, colored by original S-phase annotation (True/False). **B** Mean fraction of counts computed from histone genes as a function of inferred phase. **C** Polar histogram of inferred peak phases for cell-cycle genes, colored by their annotated peak phase according to Cyclebase [28]. **D** Boxplot displaying the variance between the mean inferred Δ*µ* of the principal cell types (*>*1000 cells) for the cell cycle genes with a mean fraction of counts *>*1e-4, depending on their peaking cell cycle phase according to **C**. **E–F** Reconstructed expression profile as a function of inferred phase, comparing the different principal cell types. Profiles were obtained by binning cells according to their inferred phase and averaging log-normalized expression (log(1 + counts per 10,000)), for *Cdca7* (E), and *Kiƒ11* (F). **G–I** Distribution of cells annotated as S phase along discrete phases obtained from Seurat Cell-Cycle Scoring (G), and continuous phases derived from the principal components 1 and 2 (H), and CoPhaser (I); the Jensen–Shannon divergence between the observed phase distribution and the original S-phase distribution is reported in each panel.

**Fig. S5.**
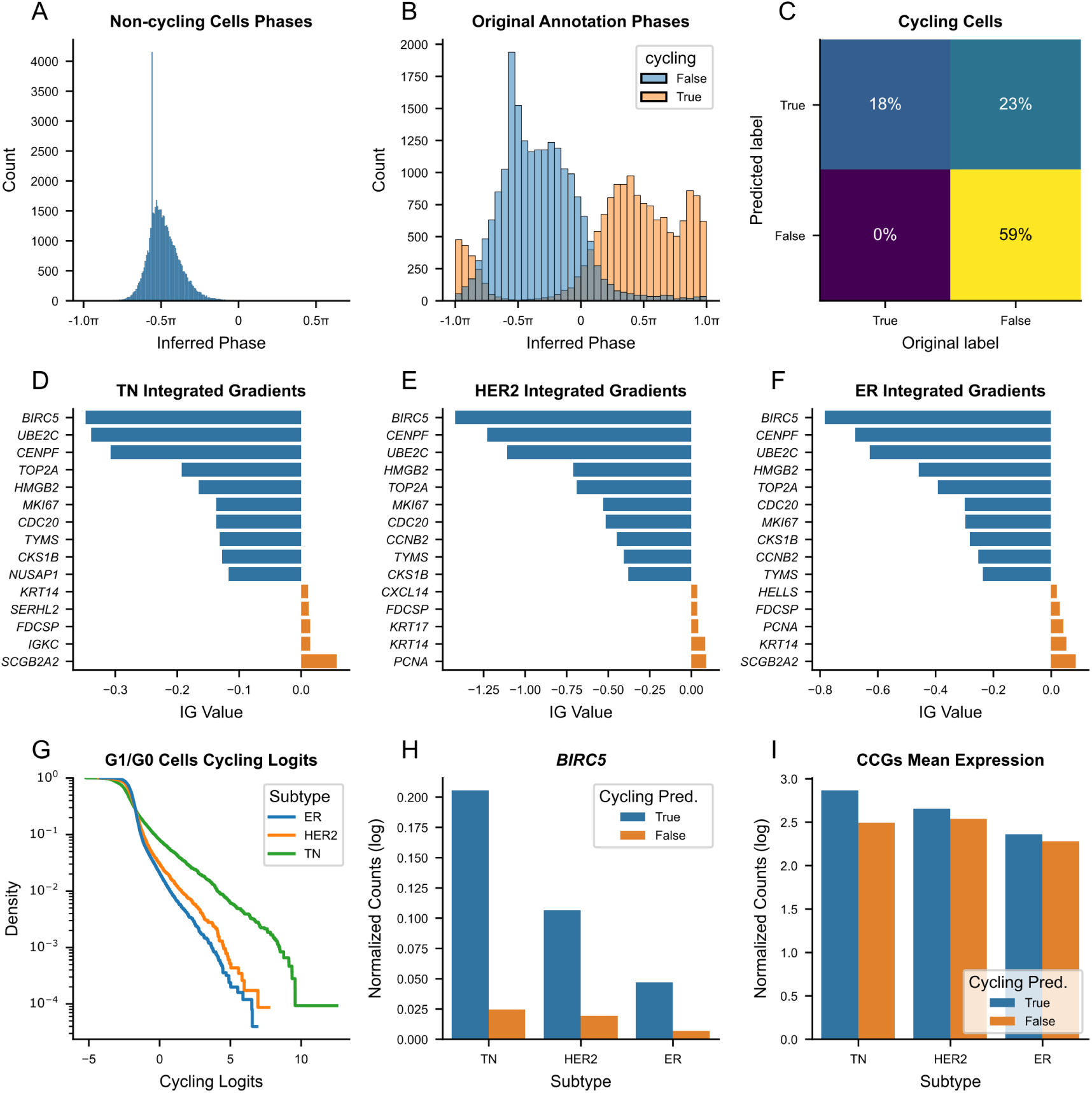
Validation of the cycling state classifier in breast cancer. **A.** Distribution of inferred phases for cells classified as non-cycling by the Bernoulli VAE. **B.** Distribution of inferred phases for cells classified as cycling by our model, colored by the original cycling status annotation. **C.** Confusion matrix comparing our model-derived cycling status with original annotations. **D–F.** Top genes contributing to the cycling classification (identified via Integrated Gradients) for three breast cancer subtypes. **G.** Empirical cumulative distribution function (ECDF) of cycling logits for cells in the G0/G1 phase, colored by subtype. **H,I.** Mean expression of *BIRC5*(**H**) and aggregate cell-cycle genes (**I**) in mid-G1 cells ([*−*0.7*π, −*0.4*π*]), comparing cells with high versus low cycling probability (weighted average) across subtypes.

**Fig. S6.**
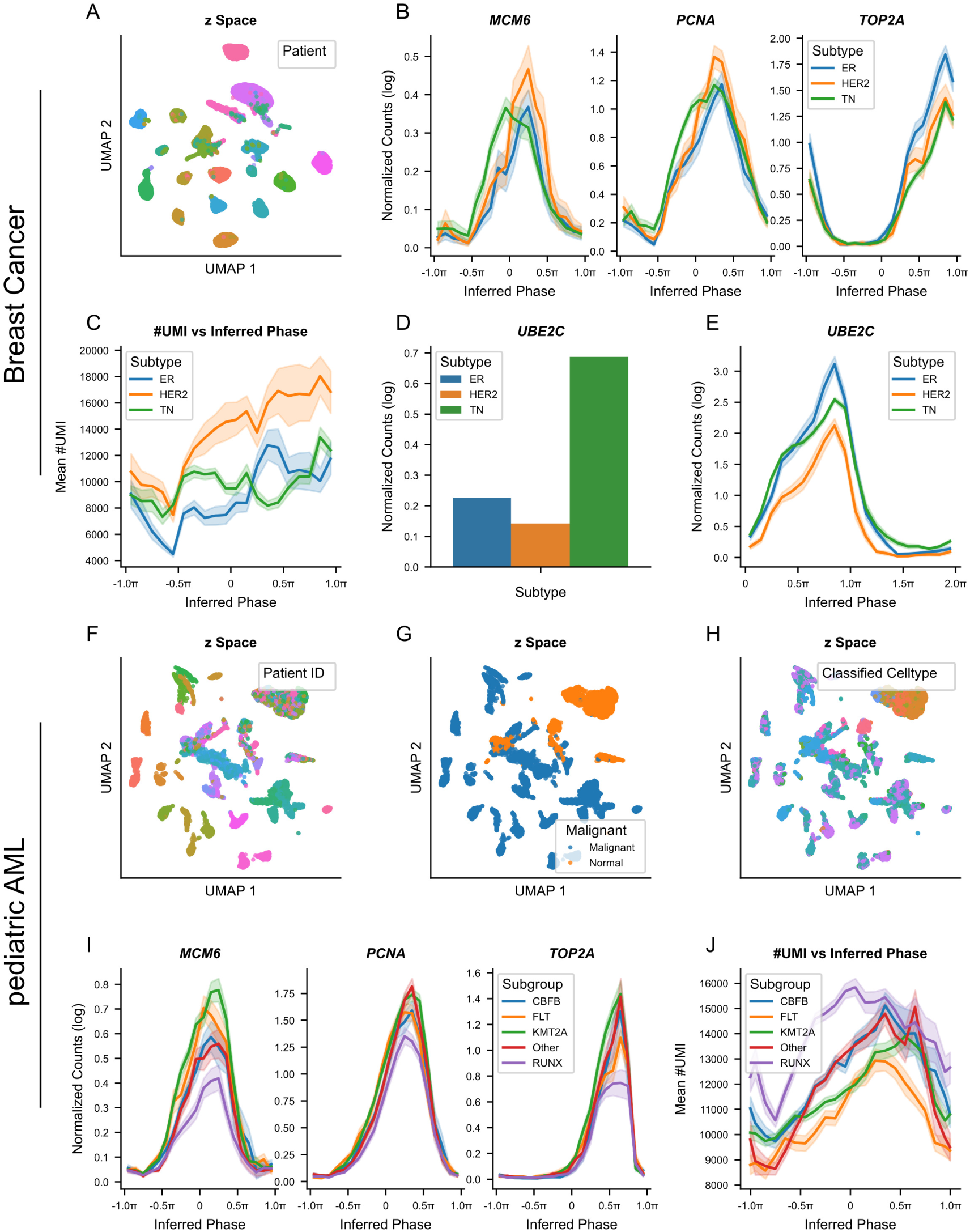
Characterization of cell-cycle dynamics in cancer. **A–E** Extended analysis of the breast cancer scRNA-seq dataset from [70]. **A** Inferred contextual *x*-space colored by patient ID. **B** Reconstructed gene expression profiles for selected cell-cycle genes (*MCM6*, *PCNA*, *TOP2A*) as a function of inferred phase, colored by cancer subtype. Profiles were obtained by binning cells according to their inferred phase and averaging log-normalized expression (log(1+counts per 10,000)). **C** Mean number of unique molecular identifiers (UMIs) per cell as a function of inferred phase, colored by cancer subtype. **D** Pseudobulk mean expression of *UBE2C* across cancer subtypes. **E** Reconstructed expression profile of *UBE2C* as a function of inferred phase, colored by cancer subtype. Profile were obtained as in **B**. **F–J** Extended analysis of the pediatric AML dataset from [35]. **F–H** Inferred contextual *x*-space colored by patient ID (**F**), malignancy status (**G**), and classified cell type (**H**). **I** Reconstructed gene expression profiles for selected cell-cycle genes (*MCM6*, *PCNA*, *TOP2A*) as a function of inferred phase, colored by tumor subtype (ER: Estrogen receptor positive, HER2: human epidermal growth factor receptor 2 positive, and TN: triple-negative). Profiles were obtained by binning cells according to their inferred phase and averaging log-normalized expression (log(1 +counts per 10,000)). **J** Mean number of unique molecular identifiers (UMIs) per cell as a function of inferred phase, colored by tumor subtype.

**Fig. S7.**
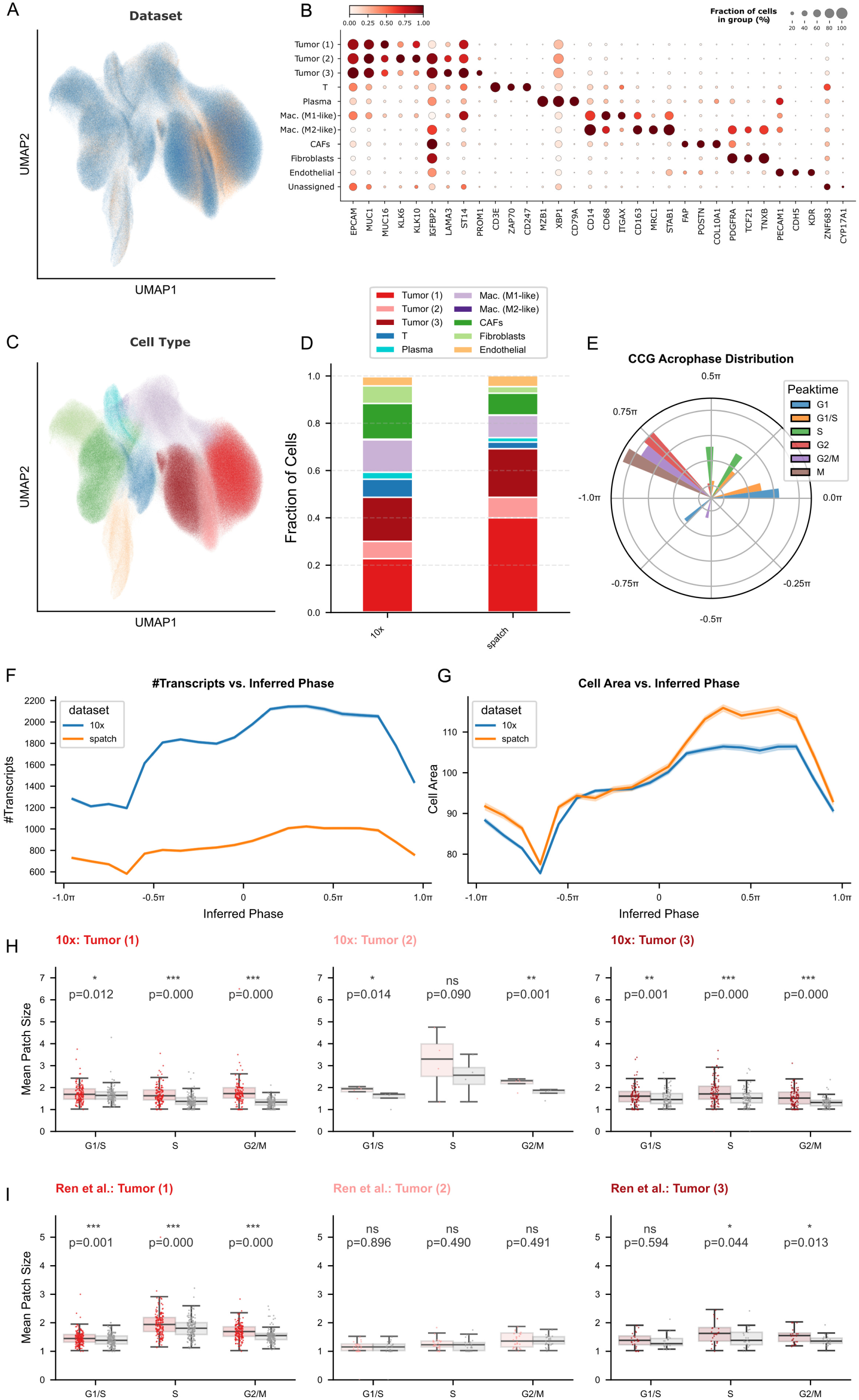
Validation of spatial cell-cycle phase inference in ovarian cancer. Xenium Prime 5K In Situ Gene Expression for two human ovarian adenocarcinoma tissues: one high-grade serous carcinoma (FFPE) [37] and one fresh frozen sample (10x Genomics). **A.** UMAP embedding of integrated z-space from both tissues, colored by dataset (10x Genomics in blue, Ren et al. in orange). **B.** Dot plot validating cell-type annotations. Dot size corresponds to the percentage of cells in the group, and color intensity represents the scaled mean expression of known marker genes. **C.** UMAP embedding of integrated z-space from both tissues, colored by cell type. **D.**. Stacked bar chart comparing cell-type composition between the 10x Genomics and Ren et al. datasets. **E.** Polar histogram of inferred peak phases for cell-cycle genes across all cell types, colored by their annotated peak phase according to Cyclebase [28]. **F.** Mean number of transcript per cell as a function of the inferred cell-cycle phase for 10x Genomics (blue) and Ren et al. (orange). **G.** Mean cell area as a function of the inferred cell-cycle phase for 10x Genomics (blue) and Ren et al. (orange). **H-I.** Spatial clustering of cell-cycle phases in tumor sub-populations for 10x Genomics (**H**) and Ren et al. (**I**). Boxplots illustrate the observed mean patch size (colored) compared to the expected null distribution (grey). The null distribution was generated via random permutations of proliferative cell-cycle phase labels (G1/S, S, G2/M) within connected components containing at least 100 cells. P-values were calculated using a two-sided paired t-test comparing observed values to null expectations across clusters (*: *p <* 0.05, **: *p <* 0.01, ***: *p <* 0.001).

**Fig. S8.**
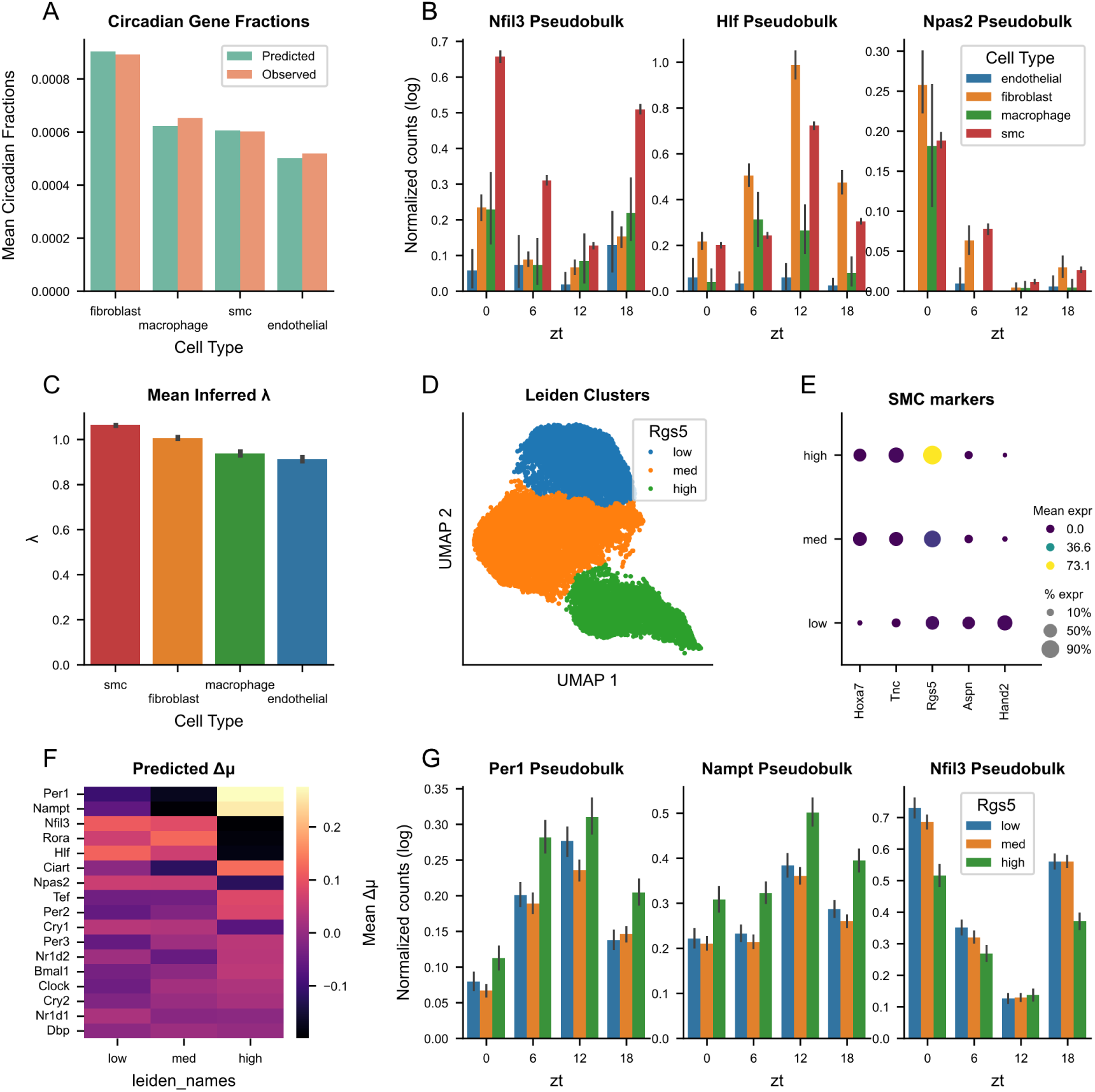
CoPhaser enables robust circadian inference in rare cell types and reveals intra-population heterogeneity. Extended analysis of mouse aorta scRNA-seq data from [21]. **A,** Comparison of the mean fraction of counts derived from circadian genes (observed) versus the model prediction, across cell types. **B,** Pseudobulk expression profiles of *Nfilh*, *Hlƒ*, and *Npas2* as a function of Zeitgeber Time (ZT), colored by cell type. **C,** Mean inferred rhythmic amplitude correction (*λ*) across cell types. **D,** Inferred contextual *x*-space for the smooth muscle cell (SMC) population, colored by Leiden cluster. **E,** Dot plot showing the expression of regional and SMC markers across the SMC leiden clusters, annotated according to their mean *Rgs5* expression. **F,** Heatmap of inferred mean shifts (Δ*µ*) for circadian rhythmic input genes across the SMC Leiden clusters. **G,** Pseudobulk expression profiles of *Per1*, *Nampt*, and *Nfilh* as a function of ZT, colored by cell type.

**Fig. S9.**
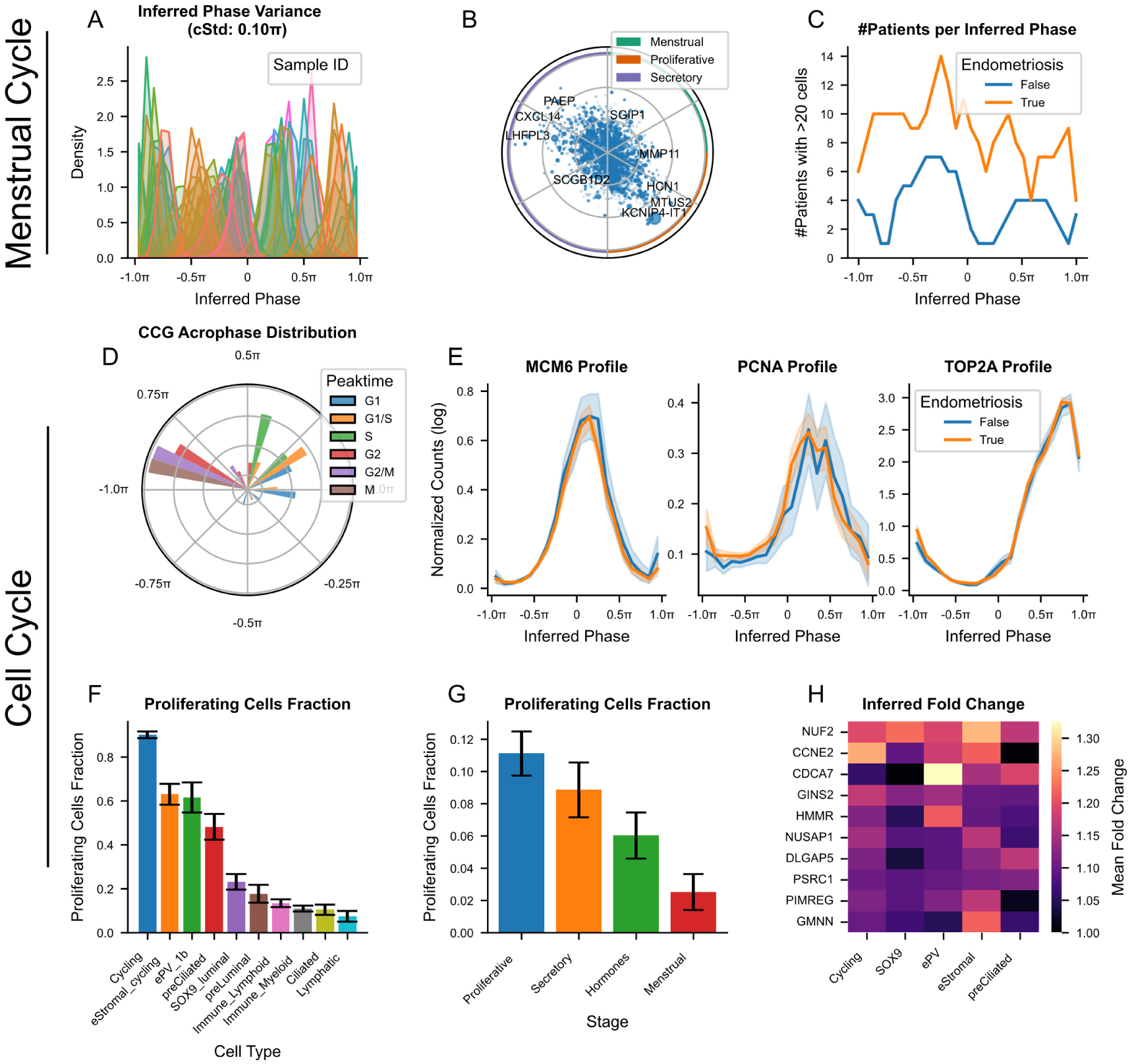
Validation of menstrual cycle reconstruction and analysis of proliferation dynamics in the human endometrium. Extended analysis of the endometrium single-nucleus RNA-seq (snRNA-seq) data from [39]. **A.** Distribution of inferred phases colored by patient ID. The mean circular standard deviation (cSTD) of phases within patients is reported (0.10*π*). **B.** Polar scatter plot of inferred amplitudes and phases for the set of context genes, showing peak-to-trough amplitude (log) as radius and peak phase as angle. Point size is proportional to mean expression. Colored rings indicate the approximate menstrual cycle stages. **C.** Number of patients with more than 20 cells in each bin of the inferred phase, comparing healthy donors and donors with endometriosis. **D.** Polar histogram of inferred peak phases for cell-cycle rhythmic input genes, from all celltypes, colored by their annotated peak phase according to Cyclebase [28]. **E.** Reconstructed expression profiles of cell-cycle genes (*MCM6*, *PCNA*, *TOP2A*) as a function of inferred cell-cycle phase, comparing cells from healthy donors and donors with endometriosis (True/False). Profiles were obtained by binning cells according to their inferred phase and averaging log-normalized expression (log(1 + counts per 10,000)). **F.** Fraction of proliferating cells across different cell types with error bars indicating the standard error of the mean (SEM) across patients. **G.** Fraction of proliferating cells across menstrual cycle stages, with error bars indicating the SEM across patients. **H.** Heatmap of the inferred mean absolute fold change in expression of cell-cycle genes between donors with endometriosis and healthy donors, across cell types.

**Fig. S10.**
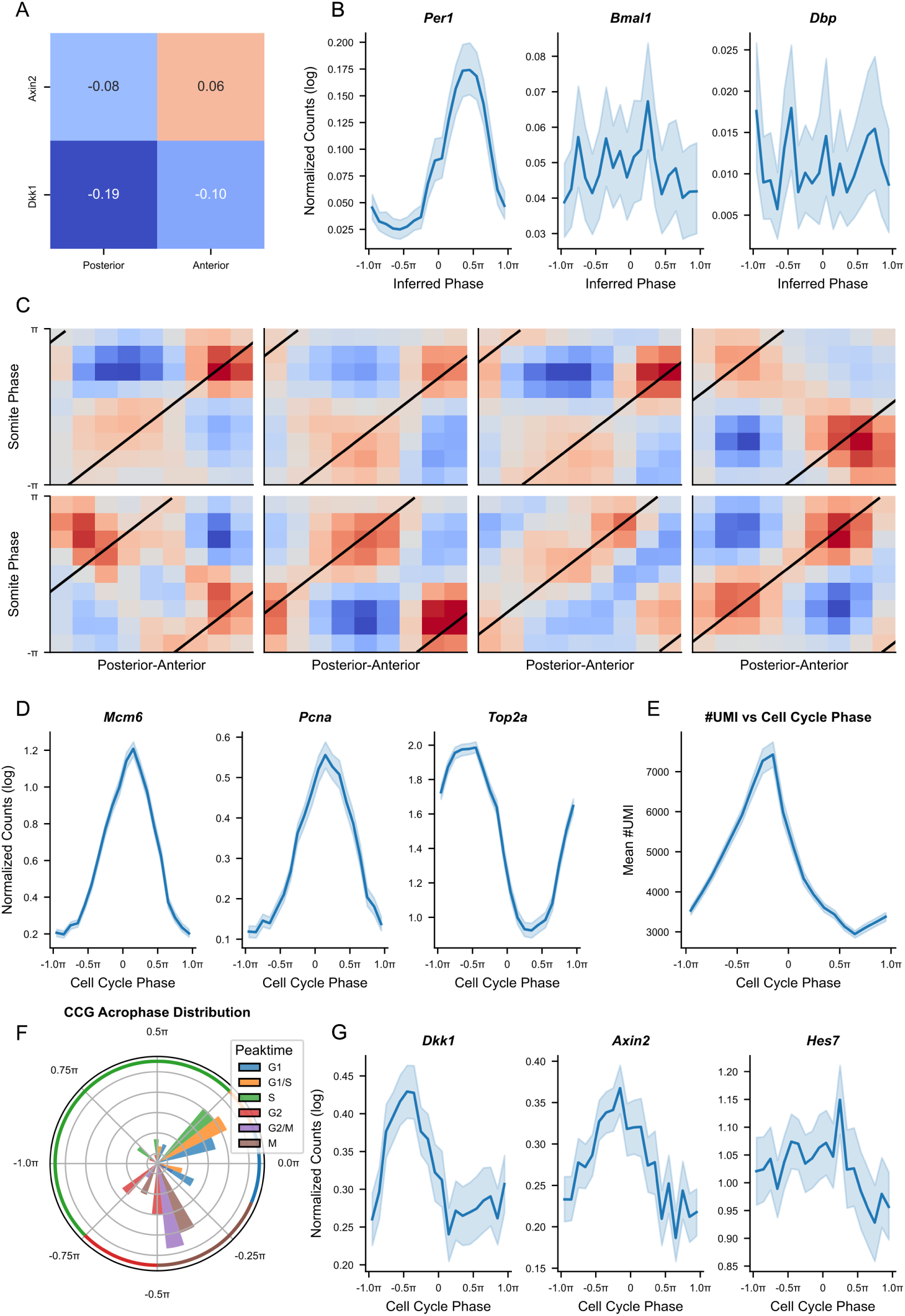
Validation of somite clock and cell cycle phase inference in the mouse PSM. Extended analysis of mouse presomitic mesoderm (PSM) data from [44]. **A.** Pearson correlation coefficients between log-normalized expression of *Axin2* and *Dkk1* and the reference gene *Hes7*, computed separately for posterior and anterior PSM cells. The posterior region corresponds to the first four quantiles of the second UMAP dimension shown in Fig. 7A, and the anterior region to the last four (out of twelve quantiles). All correlations have p-values *<* 0.001. **B.** Reconstructed expression profiles of circadian clock genes (*Per1*, *Bmal1*, *Dbp*) as a function of inferred somite clock phase. Profiles were obtained by binning cells by phase and averaging log-normalized expression (log(1 + counts per 10,000)). **C.** Representative heatmaps of pointwise mutual information (PMI) between posterior–anterior position and somite clock phase for individual embryos. PMI values range from *−*0.6 to 0.6, and embryos are ordered by their inferred traveling wave phase shift. **D.** Reconstructed expression profiles of cell cycle genes (*MCM6*, *PCNA*, *TOP2A*) as a function of inferred cell cycle phase, obtained by phase binning and averaging log-normalized expression. **E.** Mean number of unique molecular identifiers (UMIs) per cell as a function of inferred cell cycle phase. **F.** Polar histogram of inferred peak phases for rhythmic cell cycle input genes, colored by their annotated peak phase from Cyclebase [28]. The outer ring indicates the discrete cell cycle phase assignments used in Fig. 7, derived from panels **E** and **F**. **G.** Reconstructed expression profiles of somite clock genes (*Dkk1*, *Axin2*, *Hes7*) as a function of inferred cell cycle phase, obtained by binning cells by phase and averaging log-normalized expression.

## References

[2] Jovic, D., Liang, X., Zeng, H., Lin, L., Xu, F., Luo, Y.: Single-cell RNA sequencing technologies and applications: A brief overview. Clinical and Translational Medicine 12(3), 694 (2022) 10.1002/ctm2.694.

[2] Wang, L., Zhang, Q., Qin, Q., Trasanidis, N., Vinyard, M., Chen, H., Pinello, L.: Current progress and potential opportunities to infer single-cell developmental trajectory and cell fate. Current opinion in systems biology 26, 1–11 (2021)

[3] Lopez, R., Regier, J., Cole, M.B., Jordan, M.I., Yosef, N.: Deep generative modeling for single-cell transcriptomics. Nature Methods 15(12), 1053–1058 (2018)

[4] Piran, Z., Cohen, N., Hoshen, Y., Nitzan, M.: Disentanglement of single-cell data with biolord. Nature Biotechnology 42(11), 1678–1683 (2024)

[5] Lotfollahi, M., Wolf, F.A., Theis, F.J.: scGen predicts single-cell perturbation responses. Nature Methods 16(8), 715–721 (2019)

[6] Lotfollahi, M., Klimovskaia Susmelj, A., De Donno, C., Hetzel, L., Ji, Y., Ibarra, I.L., Srivatsan, S.R., Naghipourfar, M., Daza, R.M., Martin, B., Shendure, J., McFaline-Figueroa, J.L., Boyeau, P., Wolf, F.A., Yakubova, N., Günnemann, S., Trapnell, C., Lopez-Paz, D., Theis, F.J.: Predicting cellular responses to complex perturbations in high-throughput screens. Molecular Systems Biology 19(6), 11517 (2023) 10.15252/msb.202211517

[7] Rand, D.A., Raju, A., Sáez, M., Corson, F., Siggia, E.D.: Geometry of gene regulatory dynamics. Proceedings of the National Academy of Sciences 118(38), 2109729118 (2021) 10.1073/pnas.2109729118.

[8] Battich, N., Beumer, J., Barbanson, B., Krenning, L., Baron, C.S., Tanenbaum, M.E., Clevers, H., Oudenaarden, A.: Sequencing metabolically labeled transcripts in single cells reveals mRNA turnover strategies. Science 367(6482), 1151–1156 (2020)

[9] Wang, J., Symul, L., Yeung, J., Gobet, C., Sobel, J., Lück, S., Westermark, P.O., Molina, N., Naef, F.: Circadian clock-dependent and -independent posttranscriptional regulation underlies temporal mRNA accumulation in mouse liver. Proceedings of the National Academy of Sciences 115(8), 1916–1925 (2018) 10.1073/pnas.1715225115.

[10] Loureiro, C., Venzin, O.F., Oates, A.C.: Generation of patterns in the paraxial mesoderm. Current Topics in Developmental Biology 159, 372–405 (2024) 10.1016/bs.ctdb.2023.11.001

[11] Guo, X., Chen, L.: From G1 to M: a comparative study of methods for identifying cell cycle phases. Briefings in Bioinformatics 25(2), 517 (2024)

[12] Buettner, F., Natarajan, K.N., Casale, F.P., Proserpio, V., Scialdone, A., Theis, F.J., Teichmann, S.A., Marioni, J.C., Stegle, O.: Computational analysis of cell-to-cell heterogeneity in single-cell RNA-sequencing data reveals hidden subpopulations of cells. Nature Biotechnology 33(2), 155–160 (2015)

[13] Nariya, M.K., Santiago-Algarra, D., Tassy, O., Cerciat, M., Ye, T., Riba, A., Molina, N.: Single-cell multiomics reveals the oscillatory dynamics of mRNA metabolism and chromatin accessibility during the cell cycle. Cell Reports 44(8), 116089 (2025)

[14] Liang, S., Wang, F., Han, J., Chen, K.: Latent periodic process inference from single-cell RNA-seq data. Nature Communications 11(1), 1441 (2020)

[15] Riba, A., Oravecz, A., Durik, M., Jiménez, S., Alunni, V., Cerciat, M., Jung, M., Keime, C., Keyes, W.M., Molina, N.: Cell cycle gene regulation dynamics revealed by RNA velocity and deep-learning. Nature Communications 13(1), 2865 (2022)

[16] Salmen, F., De Jonghe, J., Kaminski, T.S., Alemany, A., Parada, G.E., Verity-Legg, J., Yanagida, A., Kohler, T.N., Battich, N., Brekel, F., Ellermann, A.L., Arias, A.M., Nichols, J., Hemberg, M., Hollfelder, F., Oudenaarden, A.: High-throughput total RNA sequencing in single cells using VASA-seq. Nature Biotechnology 40(12), 1780–1793 (2022)

[17] Shen, S., Vagner, S., Robert, C.: Persistent Cancer Cells: The Deadly Survivors. Cell 183(4), 860–874 (2020)

[18] Gaglia, G., Kabraji, S., Rammos, D., Dai, Y., Verma, A., Wang, S., Mills, C.E., Chung, M., Bergholz, J.S., Coy, S., Lin, J.-R., Jeselsohn, R., Metzger, O., Winer, E.P., Dillon, D.A., Zhao, J.J., Sorger, P.K., Santagata, S.: Temporal and spatial topography of cell proliferation in cancer. Nature Cell Biology 24(3), 316–326 (2022)

[19] Anafi, R.C., Francey, L.J., Hogenesch, J.B., Kim, J.: CYCLOPS reveals human transcriptional rhythms in health and disease. Proceedings of the National Academy of Sciences of the United States of America 114(20), 5312–5317 (2017)

[20] Talamanca, L., Gobet, C., Naef, F.: Sex-dimorphic and age-dependent organization of 24-hour gene expression rhythms in humans. Science 379(6631), 478–483 (2023)

[21] Auerbach, B.J., FitzGerald, G.A., Li, M.: Tempo: an unsupervised Bayesian algorithm for circadian phase inference in single-cell transcriptomics. Nature Communications 13(1), 6580 (2022)

[22] Duan, J., Ngo, M.N., Karri, S.S., Tsoi, L.C., Gudjonsson, J.E., Shahbaba, B., Lowengrub, J., Andersen, B.: tauFisher predicts circadian time from a single sample of bulk and single-cell pseudobulk transcriptomic data. Nature Communications 15(1), 3840 (2024)

[23] Xu, B., Braun, R.: VIST: variational inference for single cell time series. Genome Biology (2026)

[24] Chen, S., Regev, A., Condon, A., Ding, J.: CellUntangler: Separating distinct biological signals in single-cell data with deep generative models. Cell Genomics 0(0) (2025)

[25] Cao, N.D., Aziz, W.: The Power Spherical distribution. arXiv. arXiv:2006.04437 [stat] (2020).

[26] Belghazi, M.I., Baratin, A., Rajeshwar, S., Ozair, S., Bengio, Y., Courville, A., Hjelm, D.: Mutual Information Neural Estimation. In: Proceedings of the 35th International Conference on Machine Learning, pp. 531–540. PMLR, ???

[27] Schwabe, D., Formichetti, S., Junker, J.P., Falcke, M., Rajewsky, N.: The transcriptome dynamics of single cells during the cell cycle. Molecular Systems Biology 16(11), 20209946 (2020)

[28] Santos, A., Wernersson, R., Jensen, L.J.: Cyclebase 3.0: a multi-organism database on cell-cycle regulation and phenotypes. Nucleic Acids Research 43(Database issue), 1140–1144 (2015)

[29] Lederer, A.R., Leonardi, M., Talamanca, L., Bobrovskiy, D.M., Herrera, A., Droin, C., Khven, I., Carvalho, H.J.F., Valente, A., Dominguez Mantes, A., Mulet Arabí, P., Pinello, L., Naef, F., La Manno, G.: Statistical inference with a manifold-constrained RNA velocity model uncovers cell cycle speed modulations. Nature Methods 21(12), 2271–2286 (2024)

[30] Loaiza-Ganem, G., Cunningham, J.P.: The continuous Bernoulli: fixing a pervasive error in variational autoencoders

[31] Pal, B., Chen, Y., Vaillant, F., Capaldo, B.D., Joyce, R., Song, X., Bryant, V.L., Penington, J.S., Di Stefano, L., Tubau Ribera, N., Wilcox, S., Mann, G.B., Papenfuss, A.T., Lindeman, G.J., Smyth, G.K., Visvader, J.E.: A single-cell RNA expression atlas of normal, preneoplastic and tumorigenic states in the human breast. The EMBO Journal 40(11), 107333 (2021)

[32] Sundararajan, M., Taly, A., Yan, Q.: Axiomatic Attribution for Deep Networks. arXiv. arXiv:1703.01365 [cs] (2017).

[33] Le, X., Chen, Q., Wen, Q., Cao, S., Zhang, L., Hu, L., Hu, G., Li, Q., Chen, Z.: Design, synthesis and optimization of Apcin analogues as Cdc20 inhibitors for triple-negative breast cancer therapy. European Journal of Medicinal Chemistry 289, 117434 (2025)

[34] Verma, A., Singh, A., Singh, M.P., Nengroo, M.A., Saini, K.K., Satrusal, S.R., Khan, M.A., Chaturvedi, P., Sinha, A., Meena, S., Singh, A.K., Datta, D.: EZH2-H3K27me3 mediated KRT14 upregulation promotes TNBC peritoneal metastasis. Nature Communications 13, 7344 (2022)

[35] Lambo, S., Trinh, D.L., Ries, R.E., Jin, D., Setiadi, A., Ng, M., Leblanc, V.G., Loken, M.R., Brodersen, L.E., Dai, F., Pardo, L.M., Ma, X., Vercauteren, S.M., Meshinchi, S., Marra, M.A.: A longitudinal single-cell atlas of treatment response in pediatric AML. Cancer Cell 41(12), 2117–213512 (2023)

[36] Miller, I., Min, M., Yang, C., Tian, C., Gookin, S., Carter, D., Spence, S.L.: Ki67 is a Graded Rather than a Binary Marker of Proliferation versus Quiescence. Cell reports 24(5), 1105–11125 (2018)

[37] Ren, P., Zhang, R., Wang, Y., Zhang, P., Luo, C., Wang, S., Li, X., Zhang, Z., Zhao, Y., He, Y., Zhang, H., Li, Y., Gao, Z., Zhang, X., Zhao, Y., Liu, Z., Meng, Y., Zhang, Z., Zeng, Z.: Systematic benchmarking of high-throughput subcellular spatial transcriptomics platforms across human tumors. Nature Communications 16(1), 9232 (2025)

[38] Dobnikar, L., Taylor, A.L., Chappell, J., Oldach, P., Harman, J.L., Oerton, E., Dzierzak, E., Bennett, M.R., Spivakov, M., Jørgensen, H.F.: Disease-relevant transcriptional signatures identified in individual smooth muscle cells from healthy mouse vessels. Nature Communications 9(1), 4567 (2018)

[39] Marečková, M., Garcia-Alonso, L., Moullet, M., Lorenzi, V., Petryszak, R., Sancho-Serra, C., Oszlanczi, A., Icoresi Mazzeo, C., Wong, F.C.K., Kelava, I., Hoffman, S., Krassowski, M., Garbutt, K., Gaitskell, K., Yancheva, S., Woon, E.V., Male, V., Granne, I., Hellner, K., Mahbubani, K.T., Saeb-Parsy, K., Lotfollahi, M., Prigmore, E., Southcombe, J., Dragovic, R.A., Becker, C.M., Zondervan, K.T., Vento-Tormo, R.: An integrated single-cell reference atlas of the human endometrium. Nature Genetics 56(9), 1925–1937 (2024)

[40] Wang, W., Vilella, F., Alama, P., Moreno, I., Mignardi, M., Isakova, A., Pan, W., Simon, C., Quake, S.R.: Single-cell transcriptomic atlas of the human endometrium during the menstrual cycle. Nature Medicine 26(10), 1644–1653 (2020)

[41] Pourquié, O.: The Segmentation Clock: Converting Embryonic Time into Spatial Pattern. Science 301(5631), 328–330 (2003)

[42] Meijer, W.H.M., Sonnen, K.F.: From signalling oscillations to somite formation. Current Opinion in Systems Biology 39, 100520 (2024)

[43] Cooke, J., Zeeman, E.C.: A clock and wavefront model for control of the number of repeated structures during animal morphogenesis. Journal of Theoretical Biology 58(2), 455–476 (1976)

[44] Qiu, C., Martin, B.K., Welsh, I.C., Daza, R.M., Le, T.-M., Huang, X., Nichols, E.K., Taylor, M.L., Fulton, O., O’Day, D.R., Gomes, A.R., Ilcisin, S., Srivatsan, S., Deng, X., Disteche, C.M., Noble, W.S., Hamazaki, N., Moens, C.B., Kimelman, D., Cao, J., Schier, A.F., Spielmann, M., Murray, S.A., Trapnell, C., Shendure, J.: A single-cell time-lapse of mouse prenatal development from gastrula to birth. Nature 626(8001), 1084–1093 (2024)

[45] Matsuda, M., Yamanaka, Y., Uemura, M., Osawa, M., Saito, M.K., Nagahashi, A., Nishio, M., Guo, L., Ikegawa, S., Sakurai, S., Kihara, S., Maurissen, T.L., Nakamura, M., Matsumoto, T., Yoshitomi, H., Ikeya, M., Kawakami, N., Yamamoto, T., Woltjen, K., Ebisuya, M., Toguchida, J., Alev, C.: Recapitulating the human segmentation clock with pluripotent stem cells. Nature 580(7801), 124–129 (2020)

[46] Sonnen, K.F., Lauschke, V.M., Uraji, J., Falk, H.J., Petersen, Y., Funk, M.C., Beaupeux, M., François, P., Merten, C.A., Aulehla, A.: Modulation of Phase Shift between Wnt and Notch Signaling Oscillations Controls Mesoderm Segmentation. Cell 172(5), 1079–109012 (2018)

[47] Miao, Y., Pourquié, O.: Cellular and molecular control of vertebrate somitogenesis. Nature reviews. Molecular cell biology 25(7), 517–533 (2024)

[48] Oostrom, M.J.v., Li, Y.I., Meijer, W.H.M., Noordzij, T.E.J.C., Fountas, C., Timmers, E., Korving, J., Thomas, W.M., Simons, B.D., Sonnen, K.F.: Coupling of cell proliferation to the segmentation clock ensures robust somite scaling. bioRxiv. Pages: 2025.01.10.632257 Section: New Results (2025).

[49] Liu, J., Yang, M., Zhao, W., Zhou, X.: CCPE: cell cycle pseudotime estimation for single cell RNA-seq data. Nucleic Acids Research 50(2), 704–716 (2022)

[50] Zheng, S.C., Stein-O’Brien, G., Augustin, J.J., Slosberg, J., Carosso, G.A., Winer, B., Shin, G., Bjornsson, H.T., Goff, L.A., Hansen, K.D.: Universal prediction of cell-cycle position using transfer learning. Genome Biology 23(1), 41 (2022)

[51] Srivastava, P., Wang, T., Clark, B.Z., Yu, J., Fine, J.L., Villatoro, T.M., Carter, G.J., Brufsky, A.M., Gorantla, V.C., Huggins-Puhalla, S.L., Emens, L.A., Basili, T., Silva, E.M., Reis-Filho, J.S., Bhargava, R.: Clinical-pathologic characteristics and response to neoadjuvant chemotherapy in triple-negative low Ki-67 proliferation (TNLP) breast cancers. NPJ Breast Cancer 8, 51 (2022)

[52] Prinz, S., Hwang, E.S., Visintin, R., Amon, A.: The regulation of Cdc20 proteolysis reveals a role for the APC components Cdc23 and Cdc27 during S phase and early mitosis. Current Biology 8(13), 750–760 (1998)

[53] Min, M., Rong, Y., Tian, C., Spencer, S.L.: Temporal integration of mitogen history in mother cells controls proliferation of daughter cells. Science (New York, N.Y.) 368(6496), 1261–1265 (2020)

[54] Somer, J., Mannor, S., Alon, U.: Temporal tissue dynamics from a spatial snapshot. Nature, 1–10 (2026)

[55] Liu, Y., Sinjab, A., Min, J., Han, G., Paradiso, F., Zhang, Y., Wang, R., Pei, G., Dai, Y., Liu, Y., Cho, K.S., Dai, E., Basi, A., Burks, J.K., Rajapakshe, K.I., Chu, Y., Jiang, J., Zhang, D., Yan, X., Guerrero, P.A., Serrano, A., Li, M., Hwang, T.H., Futreal, A., Ajani, J.A., Soto, L.M.S., Jazaeri, A.A., Kadara, H., Maitra, A., Wang, L.: Conserved spatial subtypes and cellular neighborhoods of cancer-associated fibroblasts revealed by single-cell spatial multiomics. Cancer Cell 43(5), 905–9246 (2025)

[56] Zahir, N., Sun, R., Gallahan, D., Gatenby, R.A., Curtis, C.: Characterizing the ecological and evolutionary dynamics of cancer. Nature Genetics 52(8), 759–767 (2020)

[57] Smith, C.B., Vinne, V., McCartney, E., Stowie, A.C., Leise, T.L., Martin-Burgos, B., Molyneux, P.C., Garbutt, L.A., Brodsky, M.H., Davidson, A.J., Harrington, M.E., Dallmann, R., Weaver, D.R.: Cell-type specific circadian bioluminescence rhythms in Dbp reporter mice. Journal of biological rhythms 37(1), 53–77 (2022)

[58] Arora, S., Cohen, N., Hu, W., Luo, Y.: Implicit regularization in deep matrix factorization. In: Proceedings of the 33rd International Conference on Neural Information Processing Systems, pp. 7413–7424. Curran Associates Inc., Red Hook, NY, USA (2019)

[59] Hendrycks, D., Gimpel, K.: Gaussian Error Linear Units (GELUs). arXiv. arXiv:1606.08415 [cs] (2023).

[60] Jang, E., Gu, S., Poole, B.: Categorical Reparameterization with Gumbel-Softmax. arXiv. arXiv:1611.01144 [stat] (2017).

[61] Maddison, C.J., Mnih, A., Teh, Y.W.: The Concrete Distribution: A Continuous Relaxation of Discrete Random Variables. arXiv. arXiv:1611.00712 [cs] (2017).

[62] Higgins, I., Matthey, L., Pal, A., Burgess, C., Glorot, X., Botvinick, M., Mohamed, S., Lerchner, A.: -VAE: LEARNING BASIC VISUAL CONCEPTS WITH A CONSTRAINED VARIATIONAL FRAMEWORK (2017)

[65] Zheng, G.X.Y., Terry, J.M., Belgrader, P., Ryvkin, P., Bent, Z.W., Wilson, R., Ziraldo, S.B., Wheeler, T.D., McDermott, G.P., Zhu, J., Gregory, M.T., Shuga, J., Montesclaros, L., Underwood, J.G., Masquelier, D.A., Nishimura, S.Y., Schnall-Levin, M., Wyatt, P.W., Hindson, C.M., Bharadwaj, R., Wong, A., Ness, K.D., Beppu, L.W., Deeg, H.J., McFarland, C., Loeb, K.R., Valente, W.J., Ericson, N.G., Stevens, E.A., Radich, J.P., Mikkelsen, T.S., Hindson, B.J., Bielas, J.H.: Massively parallel digital transcriptional profiling of single cells. Nature Communications 8(1), 14049 (2017)

[66] La Manno, G., Soldatov, R., Zeisel, A., Braun, E., Hochgerner, H., Petukhov, V., Lidschreiber, K., Kastriti, M.E., Lönnerberg, P., Furlan, A., Fan, J., Borm, L.E., Liu, Z., Bruggen, D., Guo, J., He, X., Barker, R., Sundström, E., Castelo-Branco, G., Cramer, P., Adameyko, I., Linnarsson, S., Kharchenko, P.V.: RNA velocity of single cells. Nature 560(7719), 494–498 (2018)

[67] Tirosh, I., Izar, B., Prakadan, S.M., Wadsworth, M.H., Treacy, D., Trombetta, J.J., Rotem, A., Rodman, C., Lian, C., Murphy, G., Fallahi-Sichani, M., Dutton-Regester, K., Lin, J.-R., Cohen, O., Shah, P., Lu, D., Genshaft, A.S., Hughes, T.K., Ziegler, C.G.K., Kazer, S.W., Gaillard, A., Kolb, K.E., Villani, A.-C., Johannessen, C.M., Andreev, A.Y., Van Allen, E.M., Bertagnolli, M., Sorger, P.K., Sullivan, R.J., Flaherty, K.T., Frederick, D.T., Jané-Valbuena, J., Yoon, C.H., Rozenblatt-Rosen, O., Shalek, A.K., Regev, A., Garraway, L.A.: Dissecting the multicellular ecosystem of metastatic melanoma by single-cell RNA-seq. Science 352(6282), 189–196 (2016)

[69] Karin, J., Bornfeld, Y., Nitzan, M.: scPrisma infers, filters and enhances topological signals in single-cell data using spectral template matching. Nature Biotechnology 41(11), 1645–1654 (2023)

[70] Pal, B., Chen, Y., Vaillant, F., Capaldo, B.D., Joyce, R., Song, X., Bryant, V.L., Penington, J.S., Di Stefano, L., Tubau Ribera, N., Wilcox, S., Mann, G.B., kConFab, Papenfuss, A.T., Lindeman, G.J., Smyth, G.K., Visvader, J.E.: A single-cell RNA expression atlas of normal, preneoplastic and tumorigenic states in the human breast. The EMBO journal 40(11), 107333 (2021) 10.15252/embj.2020107333

[71] Korsunsky, I., Millard, N., Fan, J., Slowikowski, K., Zhang, F., Wei, K., Baglaenko, Y., Brenner, M., Loh, P.-r., Raychaudhuri, S.: Fast, sensitive and accurate integration of single-cell data with Harmony. Nature Methods 16(12), 1289–1296 (2019)

[72] Hu, C., Li, T., Xu, Y., Zhang, X., Li, F., Bai, J., Chen, J., Jiang, W., Yang, K., Ou, Q., Li, X., Wang, P., Zhang, Y.: CellMarker 2.0: an updated database of manually curated cell markers in human/mouse and web tools based on scRNA-seq data. Nucleic Acids Research 51(D1), 870–876 (2022)

[73] Endres, D.M., Schindelin, J.E.: A new metric for probability distributions. IEEE Transactions on Information Theory 49(7), 1858–1860 (2003)

